# DNA storage in thermoresponsive microcapsules for repeated random multiplexed data access

**DOI:** 10.1101/2023.03.17.533163

**Authors:** Bas W.A. Bögels, Bichlien H. Nguyen, David Ward, Levena Gascoigne, David P. Schrijver, Anna-Maria Makri Pistikou, Alex Joesaar, Shuo Yang, Ilja K. Voets, Willem J.M. Mulder, Andrew Phillips, Stephen Mann, Georg Seelig, Karin Strauss, Yuan-Jyue Chen, Tom F. A. de Greef

## Abstract

Owing to its longevity and extremely high information density, DNA has emerged as an attractive medium for archival data storage. Scalable parallel random access of information is a desirable property of any storage system. For DNA-based storage systems, however, this yet has to be robustly established. Here we develop thermoconfined PCR, a novel method that enables multiplexed, repeated random access of compartmentalized DNA files. Our strategy is based on stable localization of biotin-functionalized oligonucleotides inside microcapsules with temperature-dependent membrane permeability. At low temperatures, microcapsules are permeable to enzymes, primers, and amplified products, while at high temperatures membrane collapse prevents molecular crosstalk during amplification. We demonstrate that our platform outperforms non-compartmentalized DNA storage with respect to repeated random access and reducing amplification bias during multiplex PCR. Using fluorescent sorting, we additionally demonstrate sample pooling and data retrieval by barcoding of microcapsules. Our thermoresponsive microcapsule technology offers a scalable, sequence-agnostic approach for repeated random access of archival DNA files.

## Main

While the world is generating increasingly more data, our capacity to store this information is lagging behind^1^. Traditional long-term storage media such as hard disks or magnetic tape are limited in terms of durability and storage density, which has led to increased interest in, small organic molecules^2,3^, polymers^4,5^, and more recently DNA^6–8^as molecular data carriers. Owing to its intrinsic capacity for information storage, longevity and high information density, DNA in particular is a prime candidate for archival digital data storage ^9^. Significant progress has been made in coding schemes^7,10,11^ used to convert binary data to DNA, with the current best method achieving a density of 17 Eb/g^12^, exceeding magnetic and optical alternatives by approximately six orders of magnitude ^9^.

To unlock DNA’s full potential as a data storage solution three challenges in the physical domain must be overcome. First, scalable synthesis is a prerequisite for economic, widespread implementation. Current DNA-based file systems almost exclusively rely on chemically synthesized DNA of around 200 base pairs (bp). While parallel synthesis of DNA is now common^13,14^, it relies on phosphoramidite technology and its harsh reaction conditions and toxic byproducts^15,16^. To overcome this reliance on harmful chemicals, enzymatic DNA synthesis^17–19^ has emerged as a cost-effective alternative which also increases the sustainability of DNA data storage solutions^20^. Second, long-term storage is of particular interest as current magnetic data carriers^21^ typically display year half-lives of 5 to 10. Under the right conditions, DNA is extremely stable, as shown by recent success in retrieving genetic information stored in samples more than a million years old^22^. Inspired by natural fossils, researchers have developed synthetic protective shells based on glass^23,24^ and other materials^25,26^. These synthetic shells extend the half-life of DNA to hundreds of years^25,27^. Third and the focus of our study is selective and high-throughput reading of DNA-encoded data.

Next Generation Sequencing (NGS) of DNA using Illumina^28^ or nanopore^29^ sequencing has enabled high throughput reading of DNA sequences. To circumvent the need for sequencing entire datasets encoded on DNA, polymerase chain reaction (PCR)-based random access has allowed for selective retrieval of encoded data from a complex pool of DNA files^8^. However, large differences in amplification during PCR have been observed after parallel amplification of multiple DNA templates^30,31^. This PCR bias negatively affects the decoding of DNA files since successful recovery of a file requires a minimum number of reads per sequence, which is dependent on the concentration of the oligonucleotide to be read^12^. Amplification bias originates from differences in length, sequence composition, GC content and secondary structure of DNA^31^. For PCR-based retrieval of single data-encoded DNA files, the effect of this bias can be alleviated by careful primer design and encoding both physical and logical redundancies, at increased cost of DNA synthesis and sequencing^7,12,32^. Ideally, retrieval of data-encoded DNA files should not be limited to a single file. However, retrieval of multiple data-encoding DNA files from a complex DNA pool is hindered by artifact formation due to molecular crosstalk^33^. Artifact formation can be circumvented during encoding; however, this introduces severe additional constraints on sequence design^34^. Thus current strategies for retrieval of multiple DNA files are based on physically separating reactions and amplifying each individual file using multiple singleplex PCR reactions^8,35,36^. Parallel amplification in a single reaction vessel has been realized using emulsion PCR, which segregates DNA templates using water-in-oil droplets and prevents the formation of artifacts^37,38^. Although emulsion PCR has been employed in DNA data retrieval^8,34^, the elaborate workflow, destructive nature and large quantities of organic solvents makes it unattractive for large-scale adoption.

Here we introduce a new methodology termed “thermoconfined PCR”, in which the fidelity of PCR amplification is augmented by micro-reactors with temperature-dependent membrane permeabilities. Using this strategy, we implement multiplex and repeated PCR-based access of multiple DNA files from a complex DNA pool. Our method is based on the stable encapsulation of biotinylated DNA files in individual populations of thermoresponsive, semipermeable microcapsules (Figure 1a) followed by pooling of the populations. Due to the membrane’s unique thermoresponsive permeability, molecular crosstalk and the accompanying formation of artifacts at PCR temperatures is greatly reduced, allowing faithful amplification of multiple data-encoded DNA files, comparable to emulsion PCR. However, in contrast to emulsion PCR, non-biotinylated amplicons can be retrieved and separated from the original data-encoded DNA files at room temperature without destruction of the microcompartment, due to restoration of the membrane permeability at room temperature (Figure 1b).

**Figure 1.**
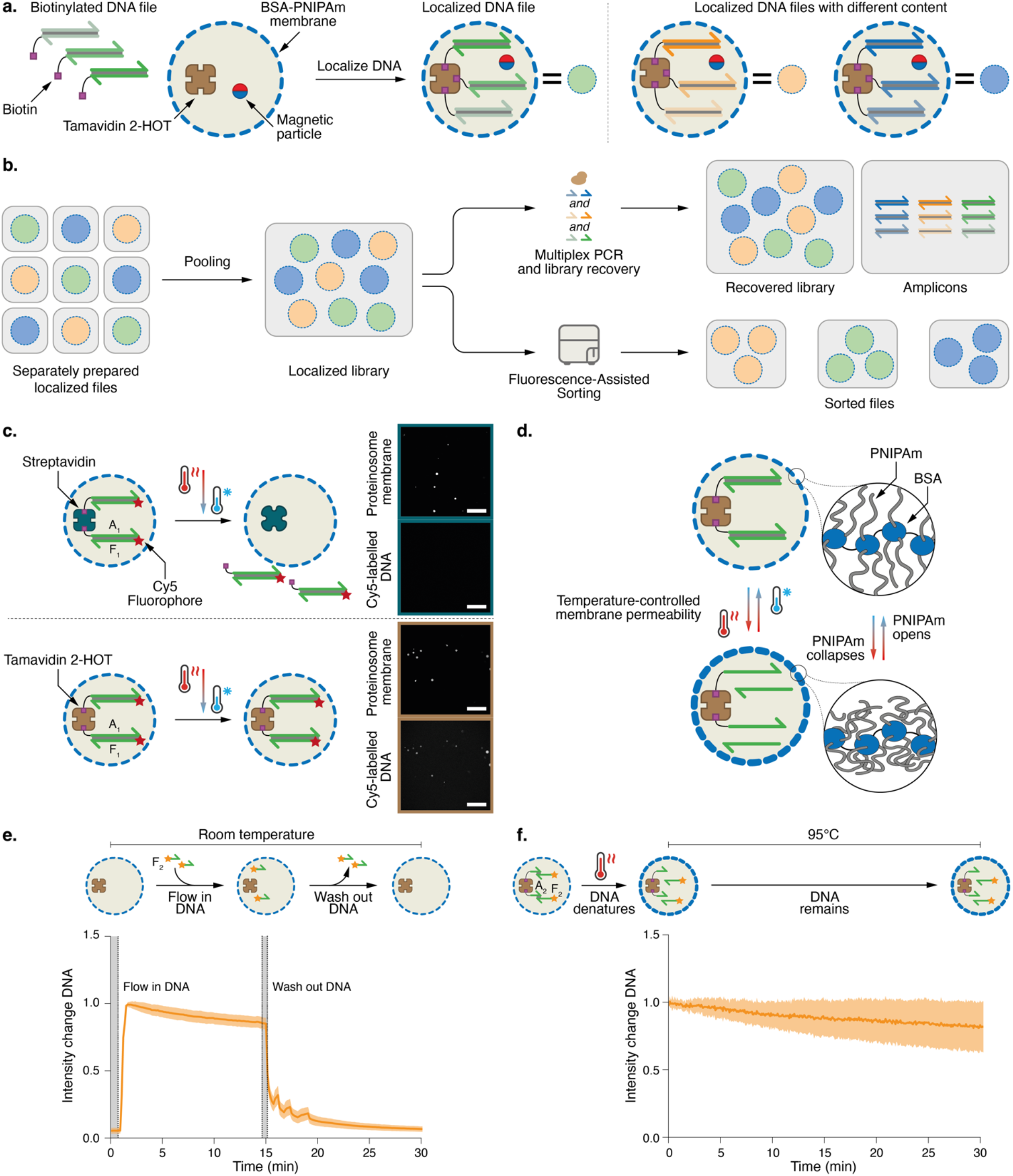
Temperature-dependent DNA localisation. **a**. Proteinosomes as a general platform for DNA data storage. Proteinosomes with BSA-PNIPAm-based thermoresponsive membranes^39^ encapsulate Tamavidin 2-HOT^45^ and magnetic particles (Methods). Digital data can be encoded into multiple DNA sequences of a fixed length which are appended with forward and reverse primer sites, these DNA templates together make up a DNA-encoded file (see Supplementary Fig. 7 for graphical representation). Using biotinylated primers during PCR, DNA files can be labelled with biotin. Biotinylated DNA files can be stably localized inside proteinosomes using the strong biotin-Tamavidin 2-HOT interaction to create a localized DNA file. Different files can be localized inside other proteinosomes to enable facile creation of multiple different localized files. **b**. After localization, files do not exchange between proteinosomes, allowing for the pooling of multiple files into a single localized library that can be amplified using multiplex PCR without molecular crosstalk and recovered using magnetic separation after PCR. Libraries can also be sorted into separate populations using fluorescence assisted sorting. **c**. Confocal micrographs of DNA containing proteinosomes after heating to 95°C and cooling. Double-stranded DNA (188 base pairs; **A**_**1**_**F**_**1**_) with Cy5 and biotin labels was localized in DyLight405-labelled proteinosomes containing 4 µM streptavidin (upper two panels) or Tamavidin 2-HOT (lower two panels) and heated to 95°C for 5 minutes and cooled to room temperature before imaging. Micrographs of the proteinosome membrane and Cy5-labelled DNA are shown in the top and bottom micrographs of each panel, respectively. dsDNA is retained only on the proteinosomes containing Tamavidin 2-HOT. Scale bar, 250 µm. The sequences for strands **A**_**1**_ and **F**_**1**_ are provided in Supplementary Table 1. **d**. Graphic showing temperature-dependent reversible closure and opening of the proteinosome membrane due to collapse and swelling of PNIPAm chains, respectively, when conjugated to crosslinked BSA (blue). High (T < LSCT) and low (T > LSCT) membrane permeabilities are shown as thin or thick blue dashed circles, respectively. **e**. Relative fluorescence intensity of fluorophore-labelled DNA inside proteinosomes at room temperature as a function of time. Proteinosomes (n=21) were trapped in a microfluidic trapping array^40^ to enable simultaneous imaging and addition or removal of reagents. A 50-nucleotide long ssDNA (**F**_**2**_) labelled with an Alexa546 fluorophore was added to the trapping array and diffusion across the membrane measured using confocal microscopy. After the fluorescent signal stabilised, buffer was added to the trapping area to remove DNA from the trapping chamber. The solid line indicates mean change, the shaded area indicates standard deviation. The sequence of strand **F**_**2**_ is provided in Supplementary Table 1. **f**. Relative fluorescence intensity of fluorophore-labelled DNA inside proteinosomes at 95°C as a function of time. A double stranded DNA complex (T_m_=65°C) consisting of 21nt biotin-labelled DNA strand (**A**_**2**_) and 50nt Alexa546 labelled strand (**F**_**2**_) was localized inside the proteinosomes (n=15). Proteinosomes are subsequently heated to 95°C while fluorescence was measured using confocal microscopy. The solid line indicates mean change, the shaded area indicates standard deviation. The sequences for strands **A**_**2**_ and **F**_**2**_ are provided in Supplementary Table 1.

Thermoconfined PCR is based on the use of proteinosomes, semipermeable microcompartments based on protein-polymer conjugates^39,40^. Biotinylated DNA files can be stably localized inside the proteinosome lumen using the encapsulated biotin-binding protein Tamavidin 2-HOT (Figure 1a). We first demonstrate heat-stable retention of biotin-labelled DNA templates inside Tamavidin 2-HOT proteinosomes. Next, we analyse the temperature dependent permeability of proteinosomes for single and double-stranded DNA using confocal fluorescence microscopy and find that the permeability of the membrane at high temperatures is significantly decreased. Triggered by this observation, we next show that molecular crosstalk during multiplex PCR amplification of two templates can be significantly reduced by isolating individual reactions inside proteinosomes. These results were then scaled to simultaneous amplification of 25 DNA files, totalling over 1.5 million unique sequences. We show that multiplex amplification of this complex DNA pool leads to a more even sequence representation when reactions are localized inside proteinosomes compared to bulk amplification. Additionally, we demonstrate that proteinosomes can enable multiple repetitive read operations by co-encapsulation of magnetic beads which allow efficient recovery of the original encapsulated DNA files after PCR-based random access. As a final step, we show that our platform is compatible with fluorescent metadata tagging using short DNA strands and membrane labels. Combined with fluorescence sorting, we show that DNA files can be selectively retrieved and amplified based on metadata, an approach that has recently been used for similarity^41^ and Boolean file search^42^.

## Results

### Heat stable DNA retention and temperature-responsive membrane permeability

We have previously^40,43,44^ shown that streptavidin-containing proteinosomes can be prepared by covalently crosslinking bovine serum albumin (BSA)/poly(N-isopropylacrylamide) (PNIPAm) conjugates at the interface of water-in-oil emulsion droplets. The crosslinked proteinosomes were phase-transferred into water and the addition of biotin-functionalized single stranded or double stranded DNA strands led to accumulation and stable retention of DNA templates up to 200bp^40^. To understand how the thermal properties of proteinosomes would influence PCR of encapsulated DNA files, we investigated the temperature-dependent stability and permeability of proteinosomes. Since streptavidin^40,43,44^ is only partially resistant to the high temperatures used during PCR we here used the heat-stable streptavidin analogue Tamavidin 2-HOT^45^. Tamavidin 2-HOT binds biotin with a similar affinity as streptavidin but can tolerate the higher temperatures during PCR. To verify the heat stability of both proteins in proteinosomes we prepared proteinosomes containing either 4µM streptavidin or 4µM Tamavidin 2-HOT (Methods, Supplementary Fig. 1) Inside these proteinosomes the 188 nucleotide (nt) long, biotinylated, Cy5-labeled double-stranded DNA (dsDNA) ***A***_***1***_***F***_***1***_ was localized by incubating the prepared proteinosomes with the dsDNA (Supplementary Fig. 2, Methods). Confocal micrographs of proteinosomes revealed that, after heating to 95°C, only proteinosomes prepared with Tamavidin 2-HOT retain internalized dsDNA (Figure 1c). To simplify downstream retrieval of proteinosomes, we also incorporated superparamagnetic particles inside the proteinosomes (see Methods). Collectively, these two changes enable heat-stable localization of DNA inside proteinosomes coupled with magnetic retrieval.

As PNIPAm becomes partially immiscible above its lower critical solution temperature (LCST, *ca*. 32°C), we reasoned that decreases in membrane permeability above the LCST could be used to retain DNA molecules produced during PCR-based processing of captured DNA files (Figure 1d). Previous work^39^ has shown that heating proteinosomes above the LCST reduces the membrane permeability for hydrophobic molecules, possibly by increasing membrane hydrophobicity and decreasing pore size. To investigate the temperature-dependent permeability for single-stranded DNA (ssDNA), we first verified that proteinosomes containing encapsulated Tamavidin 2-HOT are permeable to ssDNA at room temperature. Using a previously^40^ developed microfluidic trapping array (Supplementary Fig. 3), we captured Tamavidin 2-HOT containing proteinosomes and added a 50nt Alexa546-labeled ssDNA (***F***_*2*_) to the trapping chamber. As expected, ssDNA readily diffused across the membrane. Because ssDNA ***F***_*2*_ is not biotinylated, washing of the trapped proteinosomes results in a rapid loss of fluorescence indicative of the high permeability of the proteinosome membrane for ssDNA at room temperature (Figure 1e). Next, we prepared Tamavidin 2-HOT proteinosomes containing a biotinylated 21nt strand ***A***_***2***_ base-paired to a 10nt section of Alexa546-labeled 50nt ssDNA ***F***_***2***_ (predicted melting temperature, T_m_ = 65°C) and heated the proteinosomes to 95°C which ensures complete melting of the DNA duplex. Confocal imaging experiments at this temperature (see Methods) revealed a slow decrease of localized fluorescence over time (Figure 1f), which could be attributed to a small amount of Alexa546-labelled DNA diffusing across the membrane, or imaging artifacts (Supplementary Fig. 4), thus indicating that membrane permeability for relatively long ssDNA is much lower at this temperature. A similar experiment at 95°C using a shorter 31nt Alexa546 labelled ssDNA, similar in length to most primers, showed rapid diffusion of DNA across the membrane (Supplementary Fig. 5). Upon cooling to temperatures below the LCST, we again observe a high permeability of the membrane for both long and short ssDNA (Supplementary Fig. 6) suggesting that the temperature-induced change in membrane permeability is reversible. Together, our results reveal that Tamavidin 2-HOT proteinosomes allow stable localization of biotinylated dsDNA at high temperatures even upon melting of the DNA duplex and that the high permeability of the membrane is restored upon cooling to temperatures below the LCST.

### Enzymatic amplification of proteinosome-localized DNA

Having established that *i*) biotinylated DNA remains localized inside Tamavidin 2-HOT proteinosomes upon heating to PCR temperatures *ii)* both strands of a long duplex with a single biotin modification remain localized inside proteinosomes even upon melting of the duplex and *iii)* membrane permeability can be controlled by temperature, we proceeded to enzymatically amplify localized DNA templates. Previously^8,46^, both PCR and strand displacement amplification (SDA) have been used to amplify data-encoding DNA molecules (see Supplementary Fig. 8). To demonstrate the general applicability of proteinosomes containing internalized Tamavidin 2-HOT for DNA data retrieval, we next show that localized biotinylated DNA with a length typically used for data storage can be amplified using both PCR and SDA (Figure 2a).

**Figure 2.**
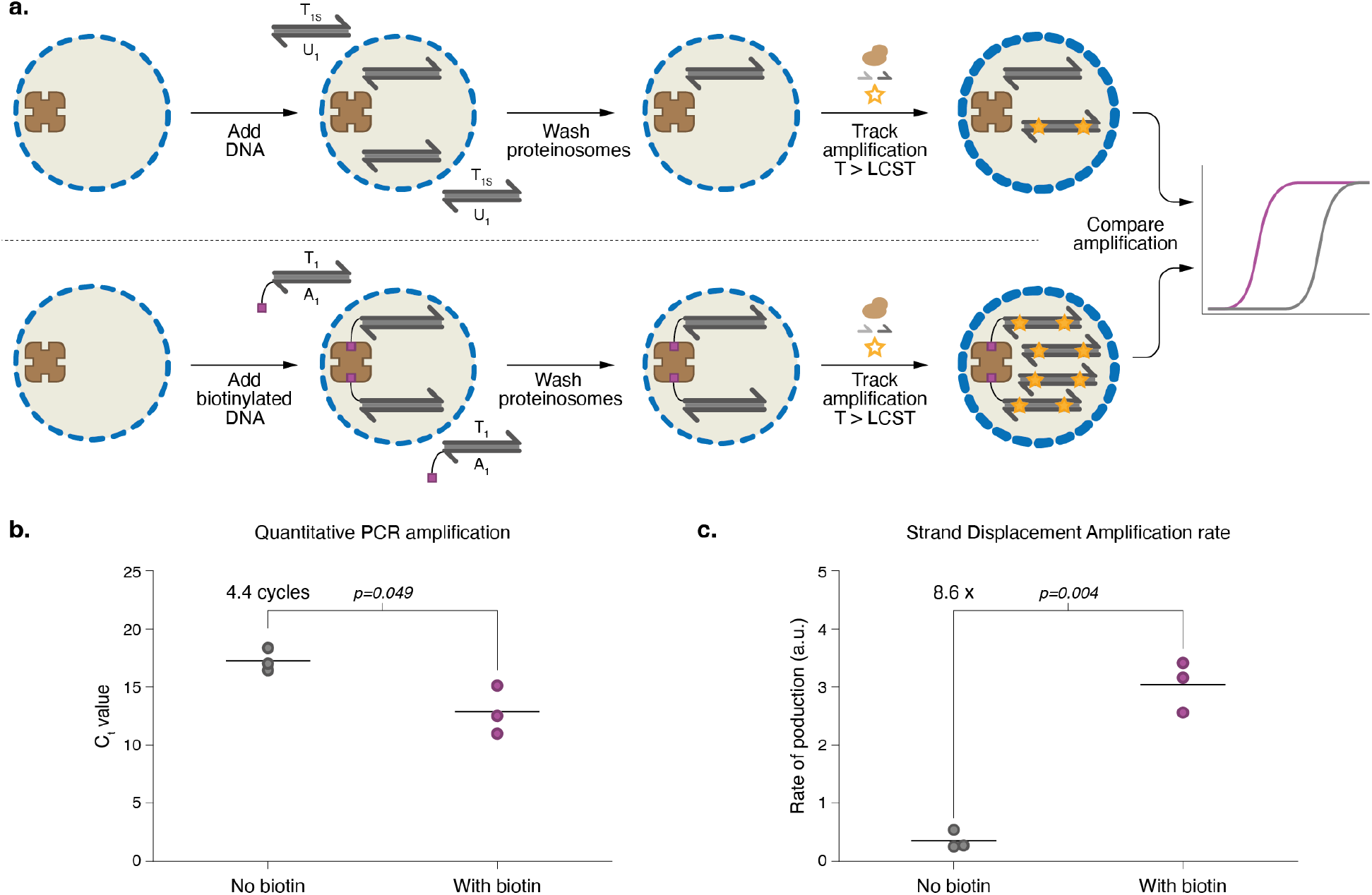
Enzymatic amplification of DNA localized inside proteinosomes. **a**. Experimental design to measure enzymatic amplification of localized DNA templates inside proteinosomes containing 4µM Tamavidin 2-HOT. DNA strands with (178 base pairs, **A**_**1**_**T**_**1**_) or without a biotin (168 base pairs, **U**_**1**_**T**_**1S**_) end modification were added to proteinosomes and removed by five washing steps (see Methods). An amplification reaction mixture containing enzymes, primers, dNTPs, and dsDNA-specific EvaGreen dye was then added and amplification of DNA localized in the proteinosomes was measured using real-time fluorescence monitoring. States of high (T < LSCT) and low (T > LSCT) membrane permeabilities are shown as thin or thick blue dashed circles, respectively. **b**. qPCR results of amplified DNA from proteinosomes incubated with either biotin-labelled DNA or non-labelled DNA (see Methods). Horizontal lines show the mean threshold cycle (C_t_) for three experiments; points represent individual experiments. A statistically significant difference of 4.4 cycles between the two conditions is observed (p=0.049). Individual amplification traces are shown in Supplementary Fig. 9. Sequences used are listed in Supplementary Table 2. **c**. SDA results of isothermally amplifying DNA from proteinosomes incubated with biotin-labelled DNA or non-labelled DNA (see Methods). Horizontal lines show the mean rate of production for three experiments; points represent individual experiments A statistically significant 8.6-fold difference in rates is observed between the two conditions (p=0.004). Individual amplification traces are shown in Supplementary Fig. 10. Sequences used are listed in Supplementary Table 2.

To measure amplification of localized dsDNA by PCR *in situ* we used quantitative PCR (qPCR, see Methods). Proteinosomes containing 4µM Tamavidin 2-HOT were incubated with 300 nM biotinylated 178nt template dsDNA ***T***_***1***_***A***_***1***_ or non-biotinylated 168nt control dsDNA ***T***_***1S***_***U***_1_. After washing to remove free template, we added a PCR reaction mixture consisting of polymerase, dNTPs, primers, and dsDNA-sensitive fluorescent dye EvaGreen. For each reaction, the threshold cycle was determined (see Methods) to assess the amount of DNA accessible for amplification. On average, 4.4 fewer cycles were needed to reach the fluorescent threshold when biotinylated dsDNA was used instead of non-biotinylated dsDNA (Figure 2b). Assuming perfect amplification during each PCR cycle, this indicates around 21 times more dsDNA is available for amplification when dsDNA is biotinylated compared to background levels, demonstrating that proteinosome-localised dsDNA is accessible for amplification by PCR.

Next, we performed isothermal amplification of localized dsDNA using SDA. As above, proteinosomes containing 4µM Tamavidin 2-HOT were incubated with 300 nM biotinylated template dsDNA ***T***_***1***_***A***_***1***_ or unlabelled ***T***_***1S***_***U***_1_ before washing and addition of amplification mixture consisting of polymerase, ssDNA binding protein, nickase, dNTPs, primers and EvaGreen. We tracked DNA production using dsDNA-sensitive EvaGreen, but since SDA is a linear process, we extracted rates of production from the fluorescence data (see Methods). SDA reactions of proteinosomes containing ***T***_***1***_***A***_***1***_ showed, on average, an 8.6 times higher rate of amplification than proteinosomes incubated with non-biotinylated ***T***_***1S***_***U***_***1***_ (Figure 2c). We attribute the difference in effect size between SDA and PCR to the more complicated mechanism and linear nature of SDA. Together, these results reveal that proteinosome-localized, biotinylated DNA templates with a length typically used for DNA data storage can be successfully retrieved by both PCR and SDA.

### Amelioration of molecular crosstalk by thermoconfined multiplex PCR

PCR-based random access allows retrieval of encoded data from complex DNA pools^8^. However, sequences that contain short regions of high similarity are susceptible to recombination during PCR^33^. Formation of such chimeric amplicons corrupt DNA files since data is inserted at incorrect positions. Additionally, the formation of chimera products leads to PCR bias in complex pools of DNA^33^. We reasoned that the temperature-controlled membrane permeability of proteinosomes should reduce chimera formation during PCR of complex DNA pools since the templates are effectively segregated at typical PCR temperatures (Figure 3a). To investigate how thermoconfined PCR from proteinosome-localised templates influences chimera formation, we designed two sets of biotinylated 178nt long dsDNA templates (***T***_***1***_***A***_***1***_ and ***T***_***2***_***A***_***3***_) that share a 31nt complementary region such that the chimera ***C***_***1***_***C***_***2***_ formed during multiplex PCR has a length of 71 base pairs (Figure 3b). The two templates ***T***_***1***_***A***_***1***_ and ***T***_***2***_***A***_***3***_ were each localized inside separate proteinosomes containing 4µM Tamavidin 2-HOT by addition of 300nM dsDNA and washed to remove excess DNA. Proteinosome populations were mixed and amplified using multiplex PCR (see Methods) and amplicons were recovered at room temperature. As a control experiment, templates were also amplified in a bulk reaction. Native PAGE analysis (Figure 3b and Methods) reveals the formation of a chimera product ***C***_***1***_***C***_***2***_ with a length of 71 base pairs in the bulk reaction. A faint band at 71bp is also observed for thermoconfined multiplex PCR, but this band has considerably lower intensity compared to the bulk PCR. Both reactions produced approximately the same amount of target DNA, as judged by the intensity of the bands at 178bp. To quantify the effect of thermoconfinement during multiplex PCR we measured the intensities of the target and chimera bands using FIJI^47^ (see Methods and Supplementary Fig. 11). We used the intensity of the target band as an internal control to account for gel-to-gel variability and calculated the ratio of chimera-to-product for each reaction (Fig. 3c). Higher ratios of chimera to target are observed for bulk reactions, indicating more chimera is formed under bulk conditions comparted to proteinosome-localized amplification. This result demonstrates thermoconfined multiplex PCR in proteinosomes significantly reduces the formation of chimeras by localizing DNA amplification to individual compartments.

**Figure 3.**
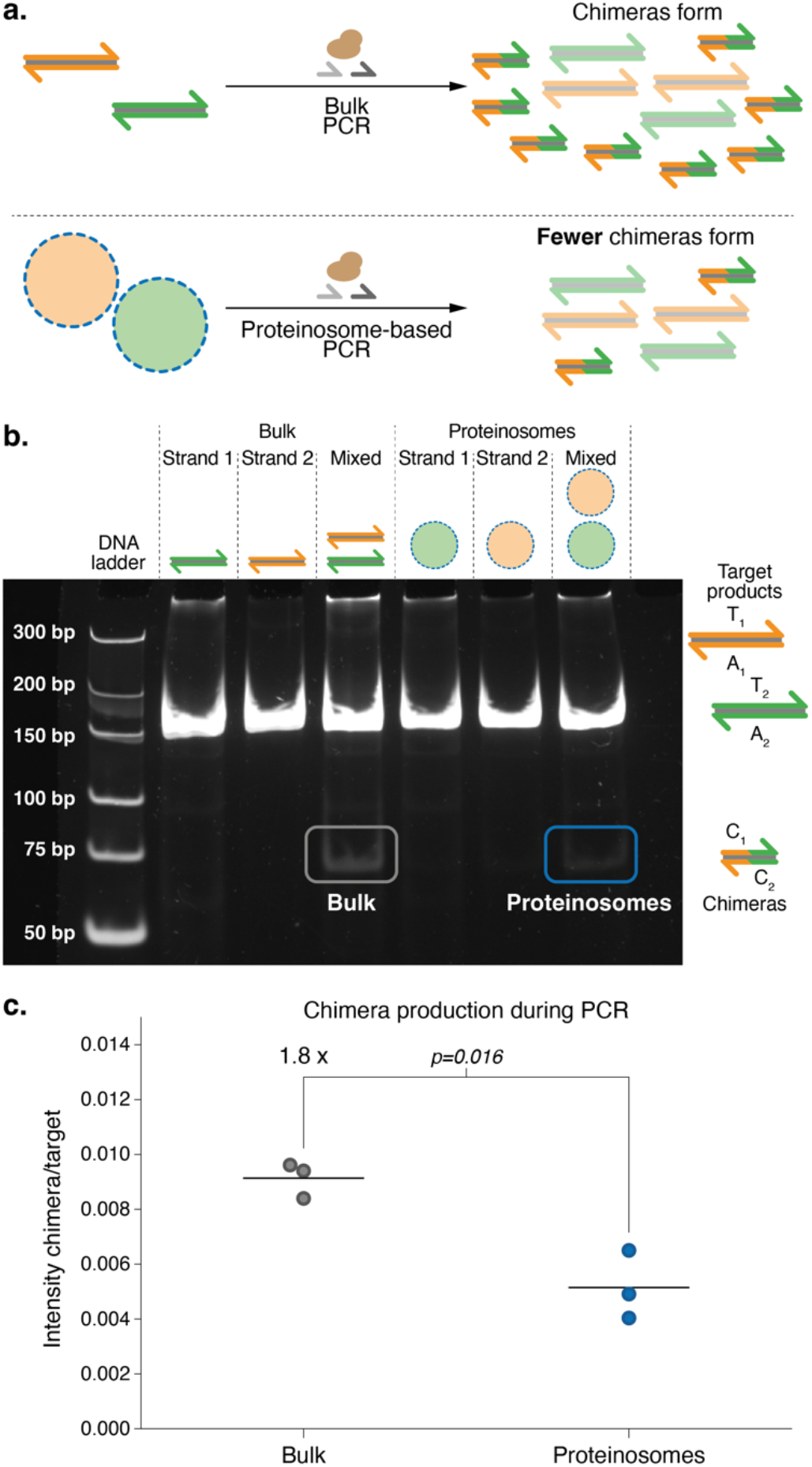
Localized amplification reduces molecular crosstalk during PCR. **a**. Thermoconfined PCR within proteinosomes reduces chimera formation during multiplex PCR by limiting molecular crosstalk. **b**. Representative native PAGE gel of reaction mixtures. Unpurified mixtures of simplex and duplex PCR in bulk or within proteinosomes were loaded on a native PAGE gel and stained using SYBR Gold before visualisation. Target strands are 178 base pairs (bp) and designed such that chimeras of 71 bp are expected to be formed. Formation of full-length 178 bp products is observed in all samples, while chimera formation is mostly observed in duplex PCR samples. Sequences used are listed in Supplementary Table 3. **c**. Quantitative analysis of PAGE results. Duplex PCR reactions were independently repeated three times and analysed using native PAGE. Intensities of target and chimera bands were determined for each reaction; the ratio between target and chimera for different methods of amplification is shown. Horizontal lines indicate the mean ratio between chimera and target intensities; data points indicate individual experiments. A statistically significant 1.8-fold difference is observed (p=0.016). Individual gel analyses are shown in Supplementary Fig 9. Sequences used are listed in Supplementary Table 3.

### Multiplex PCR and repeated access of proteinosome-localized DNA-encoded files

Having shown that spatial segregation of PCR reactions reduces chimera formation, we reasoned that localization of data-encoded files inside Tamavidin 2-HOT proteinosomes should better maintain file representation, since chimera formation is shown to negatively influence PCR bias^33^. To demonstrate this, twenty-five files stored in DNA totalling 25 MB were encoded into 110-base DNA sequences using a previously reported method^8^. Each file, consisting of approximately 66,000 unique sequences, was localized inside individual Tamavidin 2-HOT proteinosome populations before pooling 25 populations together to create a proteinosome-based library. Files were then amplified using multiplex PCR in bulk solution or compartmentalized media comprising water-in-oil emulsion droplets or proteinosome populations in water (see Methods). Relative concentrations of the files after multiplex PCR were quantified using qPCR (see Methods) to determine the fraction of each file (Supplementary Fig. 12). qPCR data revealed that thermoconfined and emulsion PCR were better able to preserve the distribution of files compared to bulk amplification. We then performed Illumina sequencing (Figure 4a) to determine the average coverage per file (see Methods and Supplementary Table 5) to test if the increased preservation translated to a more homogeneous and even coverage per file. Average coverage per file normalized to the mean was plotted on a logarithmic scale to show the deviation of coverages in orders of magnitude (Figure 4b and Methods). As expected, a large, 60-fold, difference in coverage between the most and least represented files was observed for files amplified under bulk conditions. In contrast, files that were amplified using localized reactions, showed much smaller spreads of 7- and 5-fold differences for proteinosome- and emulsion-based PCR, respectively. The spread initially present in the library prior to amplification was measured to be 3-fold. Additionally, we determined the coefficients of variation (CVs) in sequencing coverage for all conditions as a measure of evenness in the distributions. The CVs of the original pool, bulk amplified DNA, emulsion droplets and proteinosomes were 24, 139 35 and 52% respectively. These results show that thermoconfined PCR is a viable alternative to oil-based emulsions for proportional multiplex amplification of DNA encoded data.

**Figure 4.**
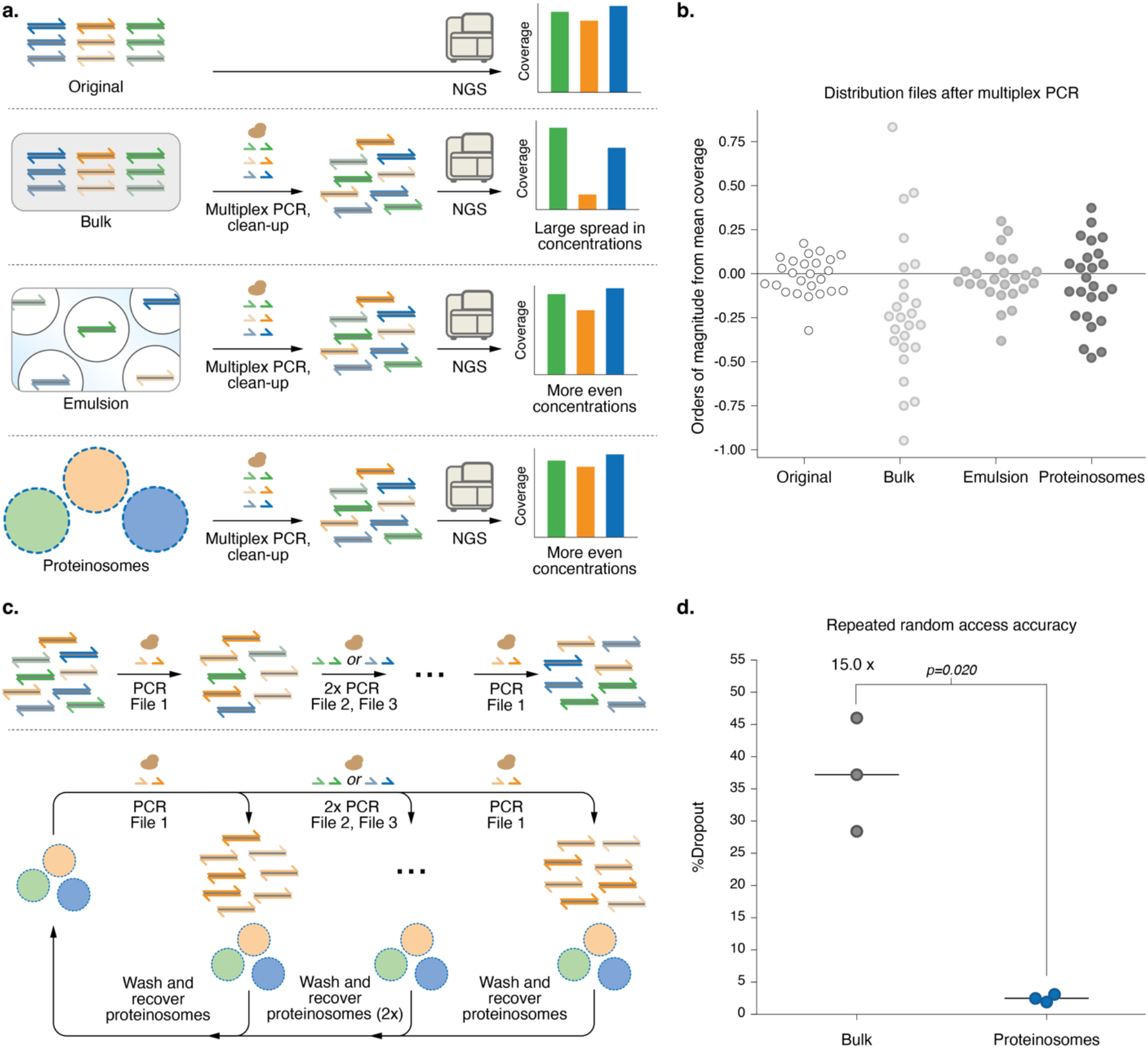
Multiplex PCR and repeated access of DNA-encoded files within proteinosomes. **a**. Schematic representation of experiments used to determine the effect of localization on DNA file concentration distributions after PCR amplification in bulk, within water-in-oil emulsion droplets or proteinosomes in water. 25 DNA-encoded files totalling 25 MB were amplified using multiplex PCR. Purified reaction mixtures were subsequently sequenced using Illumina sequencing and aligned to reference sequences to determine per file coverage (see Methods). **b** Scatterplot showing order of magnitude change from mean coverage per amplification method. Each point shows the order of magnitude deviation from the mean coverage for an individual file. 3-fold, 60-fold, 5-fold, and 7-fold changes between most and least sequenced files were observed for reference, bulk, emulsion and thermoconfined amplification, respectively. Sequences of primers used are listed in Supplementary Table 4. **c**. Schematic representation of repeated random DNA access experiment. In three consecutive PCR reactions, the files were amplified using a part of the previous reaction. After random access of the three files, the first file amplified was accessed again using PCR and the reaction mixtures were purified. Using Illumina sequencing we determined the dropout per file after the final PCR (see Methods). **d**. Dropout of sequences in the final file amplified after repeated access PCR as determined using Illumina sequencing (see Methods), randomly sampled to 30× coverage for direct comparison. Horizontal lines indicate the mean dropout of the final file accessed after four rounds of PCR for different orders of repeated random access, and points indicate individual data points. A statistically significant 15.0-fold difference in dropout between bulk- or proteinosome-based repeated random access (p=0.020) is observed. Histograms showing individual coverage distributions of the sequencing reads are shown in Supplementary Fig. 13. Sequences used are listed in Supplementary Table 4.

Since biotinylated DNA remains localized inside Tamavidin 2-HOT proteinosomes, we anticipate that recovery of the proteinosomes, and their encapsulated DNA-encoded files, using magnetic separation after PCR enables reliable repeated access of DNA encoded data. To test this, three files were localized inside proteinosomes containing 4µM Tamavidin 2-HOT that were washed before pooling the populations together to create a proteinosome-based library. Additionally, files were also mixed in equimolar concentrations in the bulk to create a non-localized library. From the resulting libraries, files were amplified in three successive rounds of PCR, each round retrieving a different file. After three rounds we again accessed the initial file (see Methods). Importantly, between PCR reactions, we recovered the proteinosome-localized library using magnetic separation (see Methods) and reused the library for amplification of the next file. In contrast, the non-localized library was PCR purified by inactivating dNTPs and primers and subsequently used for amplification of the next file (see Methods). Using Illumina sequencing we determined the dropout within a file, that is the fraction of sequences expected to be part of the file but no longer observed (Figure 4c and Methods). We observed that repeated access of the proteinosome-based library yielded a 15.0 times (p=0.020) lower dropout of sequences in the final file amplified than libraries amplified using bulk PCR (Figure 4d). Comparing the CVs of bulk and proteinosome amplified DNA also reveals that the spread is much wider for bulk amplified DNA than proteinosome-based PCR (mean bulk CV, 223%; mean proteinosome CV 73%). Together these results reveal that proteinosome-based encapsulation and subsequent pooling of large DNA libraries increases the reliability of repeated multiplex PCR operations.

### Selective file retrieval from DNA-encoded proteinosome libraries by fluorescence-assisted sorting

PCR-based retrieval of intended DNA files is very specific, however the need for sufficiently orthogonal primers limits the number of files that can be stored in a single pool^8^. This has led to development of additional strategies for random access of DNA files, such as physical separation^48^, hybridization based retrieval^41^, novel amplification schemes^36,49^ or fluorescence-assisted sorting of DNA files^42,50^. In addition to demonstrating repeated multiplex PCR, we sought to develop methods for the selective retrieval of files from DNA-encoded proteinosome libraries. To achieve this goal, we developed a strategy for fluorescent barcoding of proteinosome populations based on *i)* labelling of the microcapsule membrane with either FITC and DyLight405, and *ii)* addition of short biotinylated, Cy3 (***F***_***4***_) or Cy5 (***F***_***5***_)- labelled ssDNA after the initial data-encoded DNA files have been localized inside the proteinosomes (Figure 5a) (see Methods). This allows us to differentiate between four proteinosome populations, although in theory it is possible to generate up to 2^N^ unique barcode combinations, where *N* is the number of fluorophores used for barcoding. After encapsulation of data-encoding files in barcoded proteinosomes, we pooled the individual populations and used fluorescence activated cell sorting (FACS) to select specific files from the pool (Figure 5b) via a three-step selection procedure. First, proteinosomes were selected against unincorporated magnetic particles and background using gating on FSC-A and FSC-H channels (Supplementary Fig. 13 and 14). Second, the FITC and DyLight405 membrane labels were used as a set of fluorescence gates as the presence or absence of the fluorophores produced distinct bimodal distributions in both channels (Figure 5c left panel and Supplementary Fig. 14). Thirdly, Cy3 and Cy5 fluorescence gates were established to distinguish between the localized ***F***_***4***_ and ***F***_***5***_ sub-populations (Figure 5c right panel and Supplementary Fig. 14). Consequently, the pooled proteinosomes were sorted into four different populations based on their distinct fluorescence characteristics: (i) high DyLight405/ high Cy5; (ii) high DyLight405/high Cy3; (iii) high FITC/high Cy3; and (iv) high FITC/high Cy5. Following sorting, these four populations were measured using flow cytometry to determine the sorting accuracy (Supplementary Fig. 15). After sorting, only unimodal distributions were observed in the fluorescence data indicative of homogenous populations. Having sorted the pool into individual populations, we used qPCR to confirm that the correct DNA-encoded data was retrieved by determining the fractions of each file in the separated populations (Figure 5d). On average we found that the intended files account for 75% of all DNA in the sorted samples (non-target files were on average 8.4% each), demonstrating enrichment against other files in each sample. These results show that not only are proteinosomes compatible with PCR-based random access, but fluorescence-based metadata retrieval of proteinosomes is also possible.

**Figure 5.**
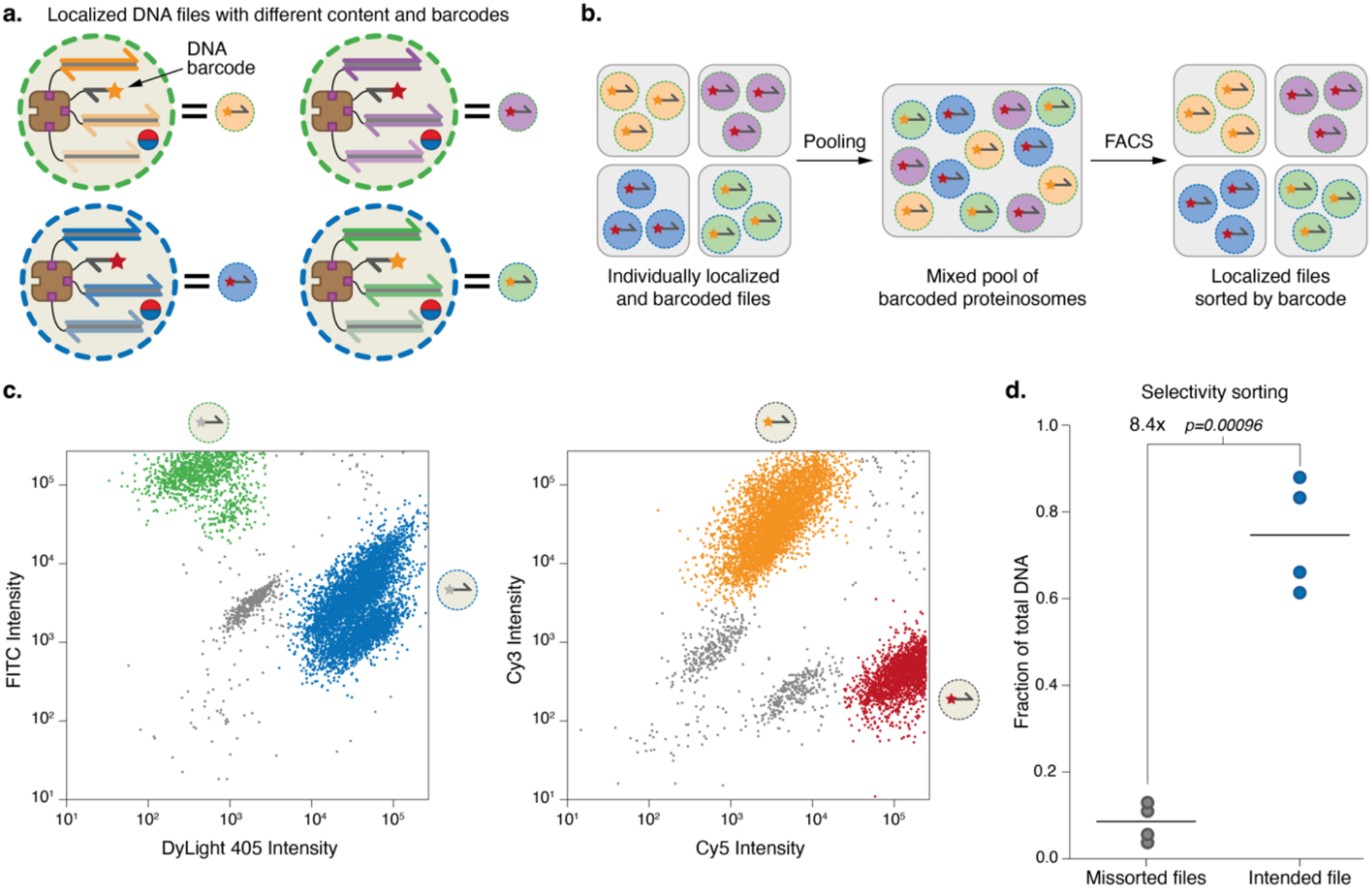
Fluorescence assisted sorting of proteinosomes for selective file retrieval. **a**. Schematic representation of membrane and localized DNA-based barcoding of proteinosomes. Proteinosome membranes are labelled with either DyLight405 (blue) or FITC (green) and additionally, fluorescent biotinylated ssDNA labelled with Cy3 (orange; **F**_**4**_) or Cy5 (red; **F**_**5**_) can be localized inside proteinosomes alongside the DNA files. **b**. Individual, barcoded populations of localized files can be pooled together in a single searchable library. Proteinosomes are sorted based on the four barcodes using FACS (see Methods). Effective sorting was verified using qPCR. **c**. FACS dot plots of FITC vs DyLight405 (left) and Cy3 vs Cy5 (right) labels. Two distinct clusters are observed in both plots which were used to sort the pooled proteinosomes into four populations. See Supplementary Fig. 14 for full gating strategy and histograms of individual fluorescent channels. Sequences for **F**_**4**_ and **F**_**5**_ are listed in Supplementary Table 5. **d**. Selectivity of sorting as determined by qPCR. Horizontal lines indicate mean fractions of files, circles indicate individual data points. Files that were unintentionally mis-sorted account for 8% of the total DNA concentration while the sorting of intended files account for 75% of the total. A statistically significant difference of 8.4-fold change in relative fractions (p=0.00096) is observed. Sequences for **F**_**4**_, **F**_**5**_ and primers used are listed in Supplementary Table 6.

## Discussion

Although NGS has significantly improved reading of DNA-encoded files, its low throughput and destructive nature of singleplex PCR random access has become a reading bottleneck in DNA data storage. In the current study, we develop proteinosomes as thermoresponsive, semipermeable microcompartments for high throughput random access of multiple DNA-encoded files using multiplex PCR. During high temperature amplification, proteinosome permeability is reduced to locally confine amplification of DNA files, thus minimizing undesired crosstalk. We demonstrate that multiplex amplification of 25 files, totalling 25 MB encoded in over 1.5 million sequences, had more proportional amplification when using proteinosomes compared to bulk PCR. In addition, we show that repeated random access can be achieved by encapsulating magnetic beads inside proteinosomes, which addresses the problem of the destructive nature of bulk and emulsion PCR. Furthermore, we show that labelling proteinosomes with fluorescent barcodes can enable fluorescence-based data retrieval. Taken together, proteinosome-based technology provides a viable path towards high-throughput random access, together with flexibility on retrieval method and repeated random access.

Although PCR- or fluorescence based random access allowed us to selectively retrieve data from a complex pool, it would still be challenging to store all data in a single pool because of slow diffusion and crosstalk, leading to inefficient random-access retrieval. Organick et al. estimated that PCR-based random access can scale up to the order of terabytes^8^. To store additional data, libraries of DNA will need to be physically separated in a compact way. While recent work has demonstrated automated microfluidic retrieval of physically separated dried spots of DNA^48^, this still requires a relatively large footprint. Using the proteinosome platform in combination with existing hard drive technology the footprint could be reduced further. Proteinosomes containing magnetic beads and DNA libraries can be physically separated and stored as individual spots on a solid support. During on demand reading, magnetic heads such as those found in hard drives could then be used to retrieve a proteinosome-DNA library for random access. The retrieved library can be subsequently recovered from the PCR mixture using magnetic heads and kept for future access. While such an approach requires further development, the present proteinosome-DNA platform provides a useful toolbox for large-scale DNA data storage. Additionally, we recently have demonstrated programmable DNA cleavage inside proteinosomes using Cas proteins^44^. Combining the broad editing capabilities of this class of proteins and the repeatable, high throughput random access described in this work would simultaneously enable editable and scalable data storage in DNA.

## Materials and methods

### Materials

2-Ethyl-1-hexanol (Sigma, 98%), 1-(3-dimethylaminopropyl)-3-ethylcarbodiimide HCl (EDC, Carbosynth), 1,6-diaminohexane (Sigma, 98%), PEG-bis(N-succinimidyl succinate) (Mw = 2000, Sigma), DyLight™ 405 NHS ester (ThermoFisher), FITC NHS ester (ThermoFisher), bovine serum albumin (heat shock fraction, pH 7, ≥98%, Sigma), streptavidin from Streptomyces avidinii (Sigma), Tamavidin 2-HOT, recombinant (Wako Chemicals), DynaBead M270 amine (Invitrogen), DynaBead MyOne carboxylic acid (Invitrogen), 1M MgCl_2_ (Invitrogen), 1M Tris pH 8.0 RNase-free (Invitrogen), 5M NaCl (Invitrogen), EvaGreen (Biotium), KAPA HiFi HotStart PCR Kit (Roche), Micellula DNA Emulsion & Purification Kit (EURx), Ibidi anti-evaporation oil (Ibidi), 30% (19:1 monomer:bis) acrylamide solution (Bio-Rad) and SYBR Gold (ThermoFisher) were used as received. All other chemicals used were purchased from Sigma. Enzymes were purchased from New England Biolabs unless noted otherwise.

### Synthesis of BSA-NH2/PNIPAm nanoconjugates

Cationized BSA (BSA-NH_2_) was synthesized according to a previously reported method^51^. Typically, a solution of diaminohexane (1.5g, 12.9 mmol in 10 ml of MilliQ water) was adjusted to pH 6.5 using 5 M HCl and added dropwise to a stirred solution of BSA (200 mg, 3 μmol in 10 ml of MilliQ water). The coupling reaction was initiated by adding 100 mg of 1-(3-dimethylaminopropyl)-3-ethylcarbodiimide HCl (EDC) immediately and then another 50 mg after 5 h. If needed, the pH value was readjusted to 6.5 and the solution was stirred for a further 6 h and then centrifuged to remove any precipitate. The supernatant was dialysed (Medicell dialysis tubing, molecular weight cutoff (MWCO) of 12−14 kDa) overnight against MilliQ water and freeze-dried.

End-capped mercaptothiazoline-activated PNIPAm (Mn = 13,284 g mol^−1^, 4 mg in 5 ml of MilliQ water) was synthesized according to the previously reported method^39^ and added to a stirred solution of BSA-NH2 (10 mg in 5 ml of PBS buffer at pH 8.0). Molecular weight and polydispersity of activated PNIPAm were determined using GPC (Supplementary Fig. 17). The solution was stirred for 10 h and then purified using a centrifugal filter (Millipore, Amicon Ultra, MWCO of 50 kDA) and freeze-dried. After freeze-drying, the obtained BSA-NH2/PNIPAm conjugate was characterized by MALDI-MS and zeta potentiometry (Supplementary Fig. 17).

FITC- and DyLight405-labelled BSA-NH_2_/PNIPAm conjugates were prepared using the same method, except that labelled BSA was used as the starting material.

### Labelling of BSA with fluorescent dyes

BSA was labelled with fluorescein isothiocyanate (FITC) as follows: 200 mg of BSA was dissolved in 10 mL of 50 mM sodium carbonate buffer (pH 9). 2.36 mg of FITC was dissolved in 590 μL of DMSO and added to the stirred BSA solution. The solution was stirred for 5 h, purified by dialyzing (Medicell dialysis tubing, MWCO 12−14 kDa) overnight against MilliQ water and freeze-dried. BSA was labelled with DyLight 405 as follows: 30 mg of BSA was dissolved in 6 mL of 50 mM sodium carbonate buffer (pH 9). 1 mg of DyLight 405 NHS ester was dissolved in 100 μL of DMF and added to the stirred BSA solution. The solution was stirred for 2 h, purified by dialyzing (Medicell dialysis tubing, MWCO 12−14 kDa) overnight against MilliQ water and freeze-dried. Using UV-Vis spectrophotometry we determined the average number of dyes per protein. For DyLight 405 labelling we measured on average 1.4 dyes per BSA molecule. For FITC labelling we measured 1.3 dyes per BSA molecule.

### Preparation of Tamavidin 2-HOT containing proteinosomes

Proteinosomes containing Tamavidin 2-HOT and magnetic particles (DynaBead M-270 amine) were prepared similarly as previously^40^ described for Streptavidin containing proteinosomes. Proteinosomes used for experiments in Fig. 1 c-e did not contain magnetic particles. BSA-PNIPAm nanoconjugates (6 mg/mL total, 1 mg/mL of which fluorescently labelled) were mixed with 4 µM Tamavidin 2-HOT and 4 mg/mL DynaBeads in 7.5 µL aqueous phase; 0.6 mg of PEG-bis(N-succinimidyl succinate) (Mw = 2,000) was dissolved in 7.5 µL 50 mM sodium carbonate buffer (pH 9) and added to the mix and briefly vortexed. Followed by the addition of 300 µL 2-Ethyl-1-hexanol and subsequent vortexing to yield the Pickering emulsion. The resulting mixture was left at room temperature for 2 hours to crosslink nanoconjugates. The oil phase was removed by pipetting away the upper oil layer and 300 µL 70% ethanol was added to resuspend the sediment. The dispersion was then sequentially dialysed (Medicell dialysis tubing, MWCO 12−14 kDa) against 70% and 50% ethanol for 2 h each and finally overnight against Milli-Q water to yield proteinosomes in water. Proteinosomes were then stored at 4°C for later use.

### DNA oligonucleotide synthesis

Except DNA encoding files from Twist Bioscience, all other DNA oligonucleotides were purchased from Integrated DNA Technologies. Modified oligonucleotides were purchased with HPLC purification, non-modified oligonucleotides were purchased desalted. Stock solutions (100 μM and 10 μM) were made using nuclease-free TE buffer (10 mM Tris, 0.1 mM EDTA, pH 7.5, Integrated DNA Technologies) and stored at −30 °C. DNA encoding for files were ordered from Twist Biosciences. These were individually PCR amplified before mixing in equal ratios in 10 ng and shipped at room temperature from Seattle to Eindhoven. These templates were then used in a similar manner as ssDNA oligonucleotides ordered from IDT.

### Preparation of double-stranded DNA

Double stranded complexes consisting of strands shorter than 100 nucleotides were formed by thermal annealing. Biotinylated strands were mixed with non-biotinylated strands at 12 and 10 µM respectively and heated to 95°C in a thermocycler for three minutes. The samples were subsequently cooled to room temperature at a rate of -0.5°C/min.

Double-stranded DNA strands longer than 100 base pairs with fluorescent and/or biotin modifications were prepared from single-stranded ultramer templates or dsDNA files and modified primers using PCR since these constructs could not be directly ordered. Typically, reactions were performed at 100 µL scale containing 5 µL (1 nM) diluted template, primers (0.5 µM each) using KAPA HiFi HotStart polymerase. The amplification protocol was as follows: 3 minutes of denaturation at 95°C, denaturation at 98°C for 20 seconds, annealing at 65°C for 15 seconds, extension at 72°C for 15 seconds. The second denaturation, annealing and extension steps were repeated 16 times followed by a final extension at 72°C for 30 seconds, before cooling down to 4°C. The resulting amplicons were then purified using Qiagen PCR extraction kit following the manufacturer’s instructions.

### DNA localization in proteinosomes

Initial localization of biotin-labelled DNA was done in 10 mM Tris (pH 8.0) with 11.5 mM MgCl_2_ and 0.1% vol/vol Tween-20. Typically, 10 µL of proteinosome containing solution was added to 5µL 4x buffer solution and 5 µL of DNA to be localized. The mixture was kept at 4°C overnight and the following day 500 µL wash buffer consisting of 10 mM Tris (pH 8.0) 1 M NaCl, 11.5 mM MgCl_2_ and 0.1% vol/vol Tween-20 was added and left at 4°C overnight and removed the following day. Secondary washing steps were performed similarly to the first except that no overnight step was used, instead proteinosomes were separated from the solution by placing the mixture in a magnetic separation rack (DynaMag, Invitrogen) for three minutes, after which the supernatant was removed by pipette.

### Initial localisation stability testing

Proteinosomes containing localized DNA were heated to 95°C in a 10µL solution using MiniPCR thermocycler for at least 5 minutes. After cooling to room temperature, a 2µL drop was then placed on a glass microscopy coverslip and confocal micrographs were taken.

### Temperature dependent Fluorescence Microscopy

Fluorescence data were acquired using a confocal laser scanning microscope (CLSM, Leica SP8) equipped with solid-state lasers (405 nm for DyLight405, 552 nm for Alexa546, 638 nm for Cy5) and a hybrid detector. The time-lapse measurements were performed with a ×10/0.40 numerical aperture (NA) (1.55 × 1.55 mm^2^ field of view, 7 μm slice thickness) at a resolution of 512 × 512 pixels. The photon-counting mode of the hybrid detector was used. High-temperature CLSM data was obtained using VAHEAT-Micro heating system (Interherence GmbH) with SmS-r substrates. A 50 µL solution of proteinosomes loaded with DNA was pipetted onto the sample cell and covered with 150 µL Ibibidi anti evaporation oil to prevent evaporation. Room temperature diffusion experiments were conducted in our previously^40^ described microfluidic trapping array. Data processing was done using custom Python code, similarly to our previously described method^40^.

### Statistics

All results reporting statistical values were from independent triplicates. Analysis was performed using Python’s SciPy (Python 3.6.5, SciPy version 1.1.0) library. In the cases where two populations were compared, Welch’s two-sided t-test was used. Statistical significance between more than two samples was determined using One-way Analysis of Variance (ANOVA) followed by post-hoc analysis, using Tukey’s multiple comparison testing. Only values of p<0.05 were considered statistically significant.

### Strand displacement amplification

Strand displacement amplification of DNA localized in proteinosomes was performed using the protocol adapted from Gao et al.^46^. Reaction mixture consisted of 1x NEB buffer 2, 0.5 µM primer, 250 µM each dNTP, 0.125 U/µL Klenow Fragment (Exo-), 0.25 U/µL Nt.BspQI, 0.2 mg/mL BSA, 4 µM T4 gene 32 protein and 1x EvaGreen. The reaction volume was 25 µL of which 2 µL consisted of proteinosomes in buffer (10 mM Tris with 11.5 mM MgCl_2_ and 0.1% vol/vol Tween-20). The reaction was kept at 37°C and recorded using a CFX96 Touch Real-Time PCR Detection System (Bio-Rad). To prevent evaporation the plate was sealed with a transparent sticker. The rate of production was determined by fitting a linear function to the fluorescence intensity and cycle number using Python.

### qPCR

Quantitative PCR was performed using CFX96 Touch Real-Time PCR Detection System (Bio-Rad). The total reaction volume was 25 µL and consisted of KAPA HiFi HotStart, 0.5 µM primers, 1x EvaGreen and 2 µL template solution (DNA or proteinosomes). Initial denaturation was set to 3 minutes at 95°C, 40 cycles of denaturation at 98°C for 20 seconds, annealing at 65°C for 15 seconds, and extension at 72°C for 15 seconds were performed followed by a final extension at 72°C for 30 seconds, before cooling down to 4°C. Fluorescence was measured during each annealing step. To prevent evaporation the plate was sealed with a transparent plate sealer. Using CFX Maestro Software (Bio-Rad) baseline correction was performed and threshold cycles (Ct) were calculated.

### Chimera formation determination using PAGE gel analysis

The total reaction volume was 25 µL and consisted of KAPA HiFi HotStart, 0.5 µM primers and 2.5 µL proteinosome solution. Thermocycling was performed in C1000 Touch thermocycler (Bio-Rad). Following initial denaturation for 3 minutes at 95°C, 20 cycles of denaturation at 98°C for 20 seconds, annealing at 65°C for 15 seconds, and extension at 72°C for 15 seconds were performed followed by a final extension at 72°C for 30 seconds, before cooling down to 4°C. To the PCR mixtures, 5 µL of 6x Loading Dye (ThermoFisher) was added before loading 12 µL on 10% TB-Mg PAGE gels. Gels were cast using 30% (19:1 monomer, bis) acrylamide solution. Running and gel buffer was 44.5 mM Tris, 44.5 mM boric acid and 11.5 mM MgCl_2_. Gels were run for 1 hour 15 minutes at 150V in Criterion vertical cell electrophoresis device (Bio-Rad) and stained for 10-15 minutes using SYBR Gold (ThermoFisher). Images were taken using an ImageQuant 400 Digital Imager (GE Healthcare). Image intensities of bands of interest were determined using ImageJ’s gel analysis plug-in. Statistical analysis was performed using Python.

### Emulsion PCR

Emulsion PCR of the library was performed using Micellula DNA Emulsion & Purification Kit. A 50µL reaction mixture containing 25 µL KAPA HiFi HotStart 2X, 50 ng template DNA, 4 µM primers (0.16 µM per pair), and 1.25 mg/mL BSA. The emulsion was formed by adding 300 µL of premixed inorganic phase and vortexing at maximum speed in a fridge at 4°C for 5 minutes, per the manufacturer’s instructions. The resultant emulsion was split into 4 tubes and thermocycled as follows: initial denaturation for 3 minutes at 95°C, 18 cycles of denaturation at 95°C for 20 seconds, annealing at 65°C for 15 seconds, and extension at 72°C for 15 seconds were performed followed by a final extension at 72°C for 30 seconds, before cooling down to 4°C. Reactions were pooled, the emulsion was broken and DNA was purified according to the manufacturer’s instructions.

The relative concentrations of individual files in the purified DNA were then quantified using qPCR.

### Multiplex PCR concentration quantification

DNA localized inside a mixed pool of proteinosomes or in bulk was amplified in 25 µL reactions consisting of 2 µL template DNA, primers (0.4 µM of each forward and reverse pair, for primer sequences used see Supplementary Table 4) using KAPA HiFi HotStart polymerase. Thermocycling was performed in C1000 Touch thermocycler (Bio-Rad). Following initial denaturation for 3 minutes at 95°C, 18 cycles of denaturation at 98°C for 20 seconds, annealing at 65°C for 15 seconds, and extension at 72°C for 15 seconds were performed followed by a final extension at 72°C for 30 seconds, before cooling down to 4°C. Reaction mixtures were purified using Qiagen PCR extraction kit following the manufacturer’s instructions.

The relative concentrations of individual files in the purified DNA were then quantified using qPCR.

### Library preparation and sequencing

Multiplex PCR reactions were performed at Eindhoven University of Technology and shipped at room temperature to University of Washington. Upon receipt, samples were validated using an Implen Nanophotometer. Subsequently, samples were prepared for sequencing following the Illumina TruSeq Nano DNA Library Prep protocol. Ends were blunted with the End-Repair buffer (ERP2), purified with Beckman Coulter AMPure XP beads, and an ‘A’ nucleotide annealed to the 3’ end with A-tailing Ligase (ATL). Ligation was performed using Illumina sequencing adapters from Illumina’s TruSeq DNA CD Indexes kit, with each sample ligated to a unique Illumina index. Finally, the samples were cleaned using Illumina Samples Purification Beads (SPB) and enriched using a 12 cycle PCR. Final product length and purity was qualified using a QIAxcel Bioanalyzer. Samples were then individually quantified using qPCR and mixed to create an equal-mass library.

A final library was prepared for sequencing by following the Illumina NextSeq Denature and Dilute Libraries Guide. Sequencing libraries were loaded in the Illumina NextSeq at 1.3 pM with a 20% control spike-in of ligated PhiX genome.

### Repeated access PCR in bulk

Three files were mixed in equal molar ratios to a final concentration of 0.5 nM. Three aliquots were amplified in 100 µL reactions consisting of 5 µL template mix, primers (0.5 µM) using KAPA HiFi HotStart polymerase. The PCR cycling protocol was initial denaturation for 3 minutes at 95°C, 10 cycles of denaturation at 98°C for 20 seconds, annealing at 65°C for 15 seconds, and extension at 72°C for 15 seconds were performed followed by a final extension at 72°C for 30 seconds, before cooling down to 4°C. A 5 µL aliquot was taken in which primers and dNTPs were then inactivated using Exo-CIP Rapid PCR Cleanup Kit (NEB). From the resulting mixture, 5 µL was then used in the next PCR as a template. The remaining 95 µL of the PCR reaction was stored at -30°C.

### Repeated access PCR in proteinosomes

Three files were localized in individual proteinosome populations washed 5 times and mixed to create the final pool. Three aliquots were amplified in 100 µL reactions consisting of 5 µL proteinosome library and primers (0.5 µM) using KAPA HiFi HotStart polymerase. The PCR cycling protocol was initial denaturation for 3 minutes at 95°C, 10 cycles of denaturation at 98°C for 20 seconds, annealing at 65°C for 15 seconds, and extension at 72°C for 15 seconds were performed followed by a final extension at 72°C for 30 seconds, before cooling down to 4°C. The reaction mixtures were then placed on a magnetic separation rack (DynaMag, Invitrogen) for three minutes to recover proteinosomes. 95 µL of the reaction mixture was pipetted off and stored at -30°C. The remaining 5 µL was washed three times using 100 µL wash buffer (10 mM Tris (pH 8.0) 1 M NaCl, 11.5 mM MgCl_2_ and 0.1% vol/vol Tween-20) and magnetic recovery of proteinosomes to the washed 5 µL solution of proteinosomes a new reaction mix was added to make a final volume of 100 µL.

### Preparation of Tamavidin 2-HOT containing proteinosomes for sorting

Due to smaller size requirements due to the nozzle used and interference of the normally used magnetic particles, proteinosomes used in the sorting experiment were prepared according to a slightly modified protocol. Proteinosomes containing Tamavidin 2-HOT and magnetic particles (DynaBeads MyOne carboxylic acid) were prepared similarly as described above. BSA-PNIPAm nanoconjugates (6 mg/mL total, 1 mg/mL of which fluorescently labelled) were mixed with 4 µM Tamavidin 2-HOT and 0.75 mg/mL DynaBeads in 20 µL total aqueous phase; 0.6 mg of PEG-bis(N-succinimidyl succinate) (Mw = 2,000) was dissolved in 40 µL 50 mM sodium carbonate buffer (pH 9) and added to the mix and briefly vortexed. Followed by the addition of 1 mL 2-Ethyl-1-hexanol and subsequent vortexing for 30 minutes to yield the Pickering emulsion. The resulting mixture was left at room temperature for 1.5 hours to crosslink nanoconjugates. The oil phase was removed by pipetting away the upper oil layer and 500 µL 70% ethanol was added to resuspend the sediment. The dispersion was then sequentially dialysed (Medicell dialysis tubing, MWCO 12−14 kDa) against 70% and 50% ethanol for 2 h each and finally overnight against Milli-Q water to yield proteinosomes in water, before filtering using a 30 µm CellTrics cell strainer (Sysmex). We verified the reduced size of these proteinosomes using confocal microscopy, the results of which are shown in Supplementary Fig. 18. Proteinosomes were then stored at 4°C for later use.

### Fluorescence assisted sorting

A FACS Aria III flow cytometer (BD Biosciences), operating at low-middle pressure, was used to interrogate a mix of proteinosomes through a 100 μm nozzle. DyLight405 and FITC labelled proteinosomes were prepared for use with FACS and files encoded in DNA were localized overnight in proteinosomes. These proteinosomes were subsequently washed with 500 µL wash buffer (10 mM Tris (pH 8.0) 1 M NaCl, 11.5 mM MgCl_2_ and 0.1% vol/vol Tween-20) and stored overnight at 4°C. The next day, 500 µL supernatant was removed and fluorescent barcode DNA (labelled with biotin and either Cy5 or Cy3) was added to proteinosomes and allowed to localize for 15 minutes at room temperature. After barcode localization proteinosomes were washed four additional times by addition of 500 µL wash buffer followed by magnetic separation and supernatant removal. Just before sorting proteinosome populations were mixed. A total of 1,000-2,000 events were recorded, from which 2D plots of the forward-scattered light height (FSC-H) versus forward-scattered light area (FSC-A), were obtained. FITC fluorescence was interrogated using a 488 nm laser and a 530/30 nm detector; Dylight405 was excited at 405 nm and detected at 460/55 nm. Cy5, ex: 633, em: 660/20. Cy3, ex: 561, em: 582/15. The gating was performed with BD FACSDiva software (BD Biosciences) and consisted of initial gating in the FSC-H vs FSC-A plot to select proteinosomes against background and unincorporated magnetic beads. From this initial gate we defined a high FITC and a high DyLight405 gate, which each were subsequently split into high Cy3 and high Cy5 gates. These were the final gates used to sort proteinosomes, for detailed gating strategy see Supplementary Fig. 15. Sorting was performed in BSA-coated tubes and sorted populations were re-analyzed using BD Aria III flow cytometer, without sorting. The final graphs were plotted using FlowCore and FlowViz packages in R.

## Acknowledgments

The authors thank P. Garvan for his assistance in running the software analysis, S. Hofstraat for help with zeta potential measurements, and G. Cremers for insightful discussions. This work was supported by Microsoft Research through its PhD Scholarship Programme, an NWO-VIDI grant from the Netherlands Organisation for Scientific Research (NWO, 723.016.003), the European Research Council (ERC project no. 101000199 AMIGA), a Vici grant from the Dutch Research Council NWO (NWO, 016.176.622).

## Author Contributions

B.W.A.B. designed the study, performed experiments, analysed the data and wrote the manuscript. L.C. and B.W.A.B. performed and analyzed high temperature microscopy experiments. D.W. and Y.C performed and analyzed the sequencing experiments. D.P.S, A.M.P. and B.W.A.B. performed and analyzed fluorescence-assisted sorting experiments. S.Y. and A.J. synthesized reagents and provided key insights during initial experiments. B.N., I.K.V., W.J.M.M., A.P, S.M., G.S, and K.S. provided critical feedback on experiments. S.M. provided critical feedback on experiments and revised the manuscript. T.F.A.d.G. and Y-J.C. conceived, designed and supervised the study, analysed the data and wrote the manuscript. All authors discussed the results and commented on the manuscript.

## Competing interests

B.N., A.P., K.S., Y-J.C. are or were employees at Microsoft Research. B.W.A.B, B.N., K.S., Y-J.C. and T.F.A.d.G. have filed patent applications on this work.

