## Supplementary Information for "DNA storage in thermoresponsive microcapsules for repeated random multiplexed data access"

### Supplementary Methods and Materials

**Preparation of fluorescent Tamavidin 2-HOT**

Fluorescent Tamavidin 2-HOT was prepared by coupling of Alexa647-NHS (Thermofisher) to Tamavidin 2-HOT. In a 20 µL reaction, 50 µM Tamavidin 2-HOT in PBS was reacted with 1 mM Alexa647-NHS dissolved in DMSO, final DMSO concentration was 12.5 v/v%. The mixture was allowed to react for 2 hours in the dark and subsequently, excess dye was removed using Zeba Micro Spin Desalting Columns, 7K MWCO (Thermofisher). Conjugate concentration was determined using NanoDrop. Using SDS-PAGE (Supplementary Fig. 1) it was determined that after purification there was still 69% unreacted dye present.

**SDS-PAGE**

For SDS-PAGE analysis 4-20% SDS-PAGE Mini-PROTEAN® TGX Precast gels (Bio-rad) were used. The running buffer consists of 25 mM Tris, 192 mM glycine, 0.1% SDS, pH 8.3. Samples were heated at 95 °C for 5 min in 1x SDS Sample Buffer (62.5 mM Tris, 10% Glycerol, 2.5% SDS (w/v), 0.01% Bromophenolblue, pH 6.8) before loading. The gels were run at 150 V for 40 minutes. Photographs of gels were taken using an ImageQuant 400 Digital Imager (GE Healthcare). Fluorescence intensities of bands of interest were determined using ImageJ's gel analysis plug-in.

**Confocal microscopy**

Quantitative fluorescence data were acquired using a confocal laser scanning microscope (CLSM, Leica SP8) equipped with solid-state lasers (405 nm for DyLight405, 488 nm for FITC, 552 nm for Alexa546, 638 nm for Cy5) and a hybrid detector (DyLight405 emission was measured at 415nm - 448nm, FITC at 498nm - 528nm, Alexa546 at 562nm - 602nm and Cy5 at 656nm - 696nm). The time-lapse measurements were performed with a ×10/0.40 numerical aperture (NA) (1.55 × 1.55 mm2 field of view, 7 µm slice thickness) at a resolution of 512 × 512 pixels. The photon-counting mode of the hybrid detector was used. High-temperature CLSM data was obtained using VAHEAT-Micro heating system (Interherence GmbH) with SmS-r substrates. A 50 µL sample solution was pipetted onto the sample cell and covered with 150 µL Ibibidi anti evaporation oil to prevent evaporation. Time course experiments at room temperature were conducted in our previously described microfluidic trapping array[1], data processing was done using custom Python code, similarly to our previously described method[1]. Single confocal micrographs were made by placing a 1 µL droplet of sample on a glass cover slide.

**Brightfield and epifluorescence microscopy**

All brightfield and epifluorescence microscopy was performed at room temperature. Imaging was performed using a Leica SP8 microscope using a ×10/0.40 numerical aperture (NA). An external light source (Leica) was used for epifluorescence imaging. To visualize DyLight405 fluorescence the DAPI fluorcube was used, FITC fluorescence was detected using FITC fluorcube. Images were captured using the attached digital camera on a field of view 1.2421x0.9313 $mm^2$ and a resolution of 960x720 pixels.

**Determining Tamavidin 2-HOT loading capacity**

The encapsulation efficiency of Tamavidin 2-HOT inside proteinosomes was determined by preparing proteinosomes as normal with AF647-labelled Tamavidin 2-HOT. These proteinosomes were imaged using confocal microscopy and the internal concentration of AF647-labelled Tamavidin 2-HOT was determined by comparing fluorescence intensity against a calibration series. Any non-reacted dye was assumed to have been removed during overnight dialysis of proteinosomes but was present in the calibration series, which we corrected for this using previously measured labelling efficiency. To prepare a calibration curve, 1 µL drops of known concentrations AF647-Tamavidin 2-HOT were placed on a glass slide and three confocal micrographs were taken at different Z-positions. Mean fluorescence intensities were determined using LASX software (Leica GmbH).

**Determining DNA loading capacity Tamavidin 2-HOT proteinosomes**

Loading capacity for DNA inside proteinosomes was determined by incubating proteinosomes containing 4µM Tamavidin 2-HOT with 300 nM biotin- and Cy5 labelled dsDNA in 10 mM Tris (pH 8.0) with 11.5 mM $MgCl_2$ and 0.1% vol/vol Tween-20. 10 µL of proteinosome containing solution was added to 5µL 4x buffer solution and 5 µL of DNA to be localized. The mixture was kept at 4°C overnight and the following day 500 µL wash buffer consisting of 10 mM Tris (pH 8.0) 1 M NaCl, 11.5 mM $MgCl_2$ and 0.1% vol/vol Tween-20 was added and left at 4°C overnight and removed the following day. Four secondary washing steps were performed similarly to the first except that no overnight step was used, instead proteinosomes were separated from the solution by placing the mixture in a magnetic separation rack (DynaMag, Invitrogen) for three minutes, after which the supernatant was removed by pipette. These proteinosomes were imaged using confocal microscopy and the internal concentration of Cy5-labelled DNA was determined by comparing fluorescence intensity against a calibration series. To prepare a calibration curve, 1 µL drops of biotin- and Cy5-labelled DNA of known concentrations were placed on a glass slide and three confocal micrographs were taken at different Z-positions. Mean fluorescence intensities were determined using LASX software (Leica GmbH).

**Gel Permeation Chromatography**

Polymer molecular weight distributions were measured using mixed-C and mixed-D columns (Polymer Labs, 300 x 70 mm) connected in series at 40°C using a Shimadzu Prominence-i system. The eluent was tetrahydrofuran (THF) and the concentration of polymer was 1.0 mg/mL. The data were processed using Shimadzu GPC software.

**MALDI-MS**

MALDI-TOF MS was performed on an Autoflex Speed instrument (Bruker, Bremen, Germany) using 2,5-dihydroxybenzoic acid as matrix substance. The sample aqueous solution concentration was 1.0 mg/mL.

**Zeta potentiometry**

Zeta potentiometry measurements were done using a Malvern ZetaSeizer Nano ZSP. Samples (0.1 mg/ml) were measured in PBS (pH 7.4 0.1 mM) buffer at room temperature. Data were analysed using Malvern ZetaSeizer software and plotted using Python.

**Characterisation size of Tamavidin 2-HOT containing proteinosomes and emulsion droplets**

The size distribution of proteinosomes and emulsion droplets was determined using fluorescence and brightfield microscopy. A 2 µL droplet containing proteinosomes or emulsion

PCR droplets was placed on a glass coverslip and a micrograph was taken. The diameter per proteinosome was determined using FIJI/ImageJ, the resulting distributions are shown in Supplementary Fig. 19.

### Supplementary Figures

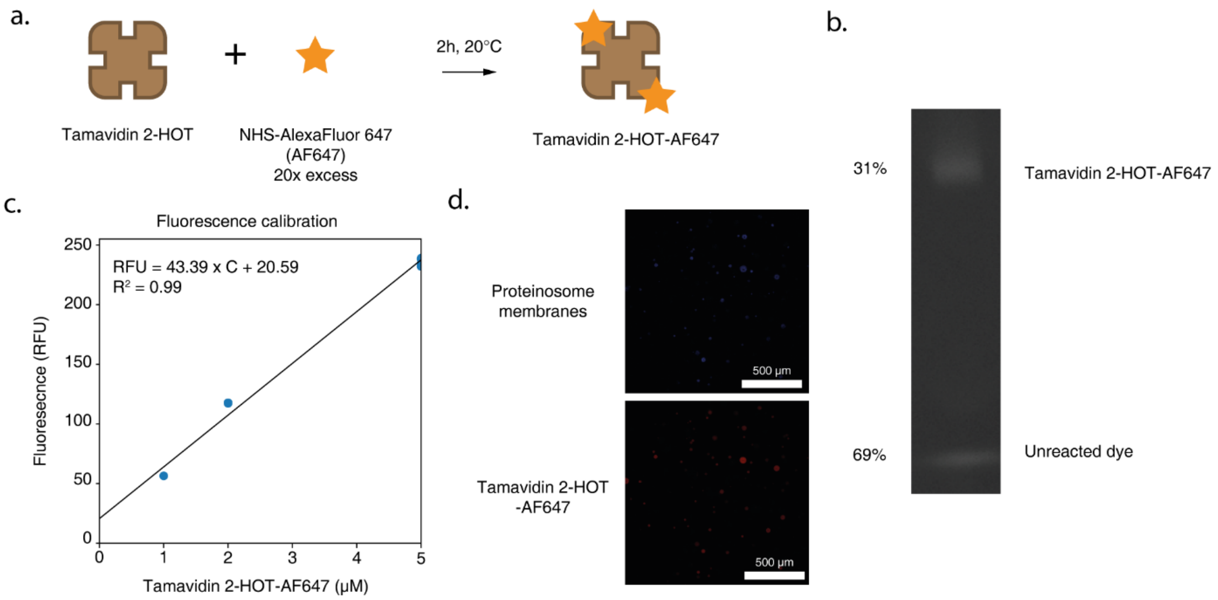

**Supplementary Figure 1 Measuring Tamavidin 2-HOT loading. a**. Fluorescent labelling strategy using NHS-AlexaFluor 647. **b.** Image of a non-reduced SDS-PAGE gel (4-20% gradient polyacrylamide, unstained) of the purified AF647-Tamavidin 2-HOT. Gel band intensity analysis was used to determine contamination of free AF647 (see Supplementary Methods). **c.** Calibration curve of fluorescence intensity vs concentration Tamavidin 2-HOT-AF647 determined at room temperature. **d.** Confocal micrographs of proteinosomes made using Tamavidin 2-HOT-AF647 at room temperature. Proteinosomes were prepared as normal, except fluorescently labelled Tamavidin 2-HOT was used. The final internal Tamavidin 2-HOT-AF657 concentration was determined using the calibration curve and accounted for the removal of unreacted dye by dividing by 0.31. The final concentration was measured to be 4.00±0.68 μM (*n=5*).

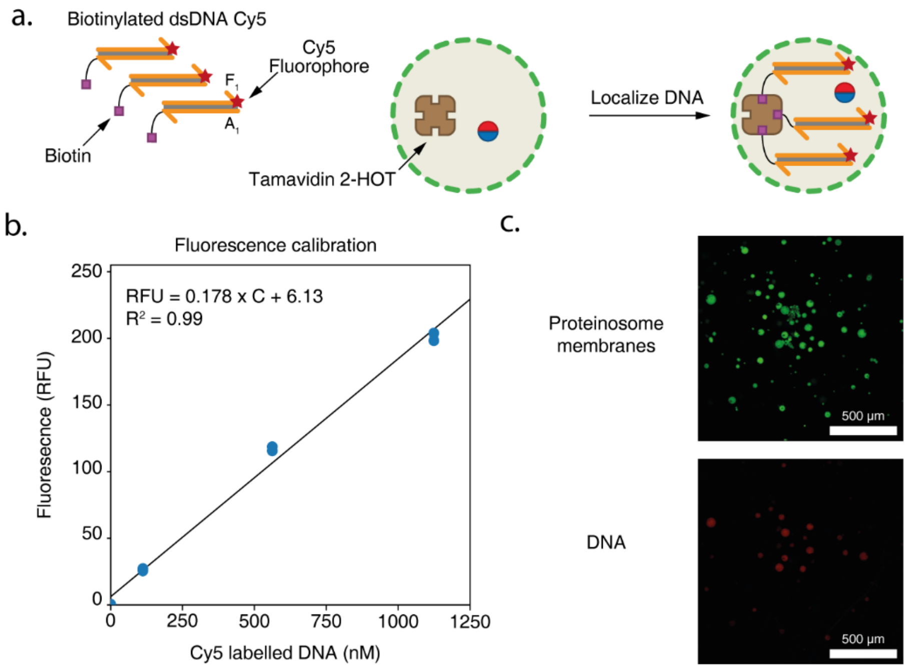

**Supplementary Figure 2 Determination dsDNA loading capacity. a.** Loading of fluorescent DNA inside Tamavidin 2-HOT proteinosomes. A 188 basepair (bp) dsDNA probe $\boldsymbol{A_1F_1}$ containing both a biotin and Cy5 label was prepared and localized inside FITC-labelled proteinosomes containing 4µM Tamavidin 2-HOT by incubating 300 nM DNA with proteinosomes overnight before washing and imaged using confocal microscopy at room temperature. **b.** Calibration curve of fluorescence intensity vs concentration Cy5- and biotin-labelled dsDNA determined at room temperature. **c.** Confocal micrographs of proteinosomes (green) made with 4µM Tamavidin 2-HOT containing localised Cy5- and biotin-labelled dsDNA (red). Fluorescence intensity of proteinosome subsections were determined and internalized dsDNA concentration was determined using a calibration curve. The final concentration of DNA was determined to be 93.8±12.7nM (*n=10*). Sequences for $\boldsymbol{A_1}$ and $\boldsymbol{F_1}$ are listed in Supplementary Table 1.

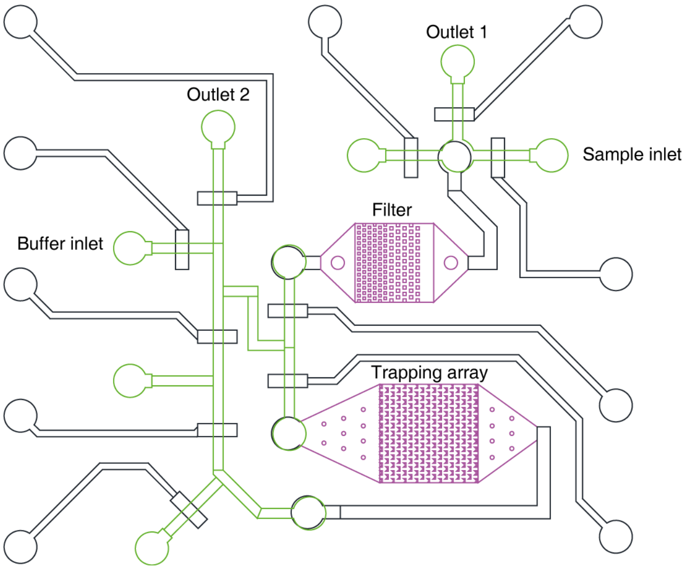

**Supplementary Figure 3 Microfluidic trapping array design.** CAD drawing of the microfluidic device, adapted from previously developed design[1,2]. Trapping array and filter are highlighted in purple. The channels of the bottom layer (bonded to the glass slide) are shown in black. The rounded top layer channels are shown in green. The flow channels crosses from the top layer to the bottom layer, as only the rounded channels of the top layer can be closed using pneumatic valves, while the trapping array requires the rectangular channels of the bottom layer. In a typical experiment proteinosomes or DNA are loaded via sample inlet port and flown through outlet 1 until no air bubbles were observed. Flow was then directed through the filter and trapping array to the outlet 2. If needed buffer could be added directly to the trapping array using the buffer inlet port.

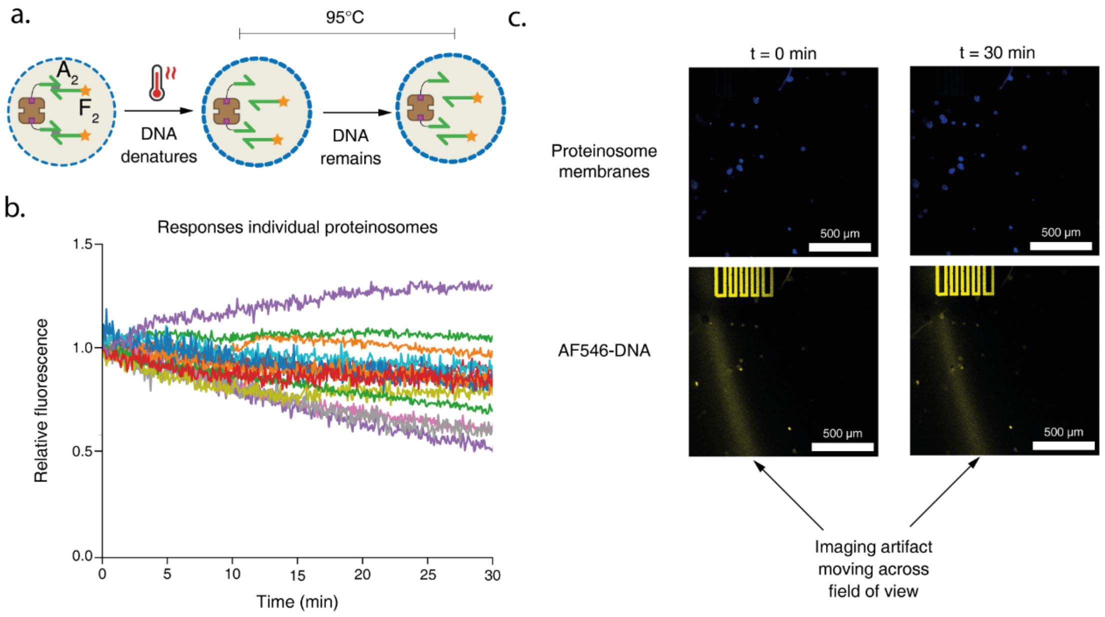

**Supplementary figure 4 Detailed images belonging to figure 1f. a.** Cartoon depicting experimental setup, as described in main text Figure 1f. **b.** Relative fluorescence intensity of fluorophore-labelled DNA inside proteinosomes at 95˚C as a function of time. A double stranded DNA complex ($T_m$=65°C) consisting of 21 nt biotin-labelled DNA strand (***$A_2$***) and 50 nt Alexa546 labelled strand (***$F_2$***) was localized inside proteinosomes (n=15). Proteinosomes are subsequently heated to 95°C while Alexa546 fluorescence was measured using confocal microscopy. Each trace is the mean fluorescence of an individual proteinosome relative to its fluorescence at t = 0 minutes. **c.** Confocal micrographs of proteinosomes containing AF546-labelled DNA in VAHEAT heating slides at times t=0 minutes and t=30 minutes. A reflection is observed to move across the imaging field of view in the AF546-DNA channel, complicating imaging. Sequences for ***$A_2$*** and ***$F_2$*** are listed in Supplementary Table 1.

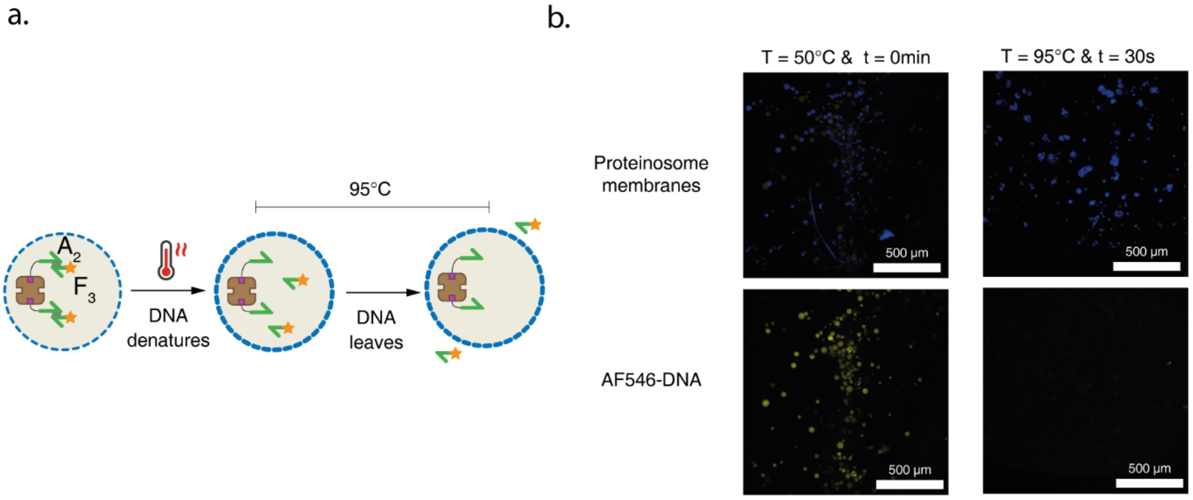

**Supplementary figure 5 Diffusion of 31mer ssDNA over proteinosome membranes is not affected by temperature. a.** Cartoon depicting experimental setup. A double stranded DNA complex (melting temperature 65°C) consisting of 21nt biotin-labelled DNA strand (A2) and 31nt Alexa546 labelled strand (***$F_3$***) was localized inside proteinosomes. Proteinosomes are subsequently heated to 95°C and subsequently imaged using confocal microscopy. **b.** Confocal micrographs t=0 minutes and 50°C prior to heating and t=30 seconds and 95°C. Sequences for ***$A_2$*** and ***$F_3$*** are listed in Supplementary Table 1.

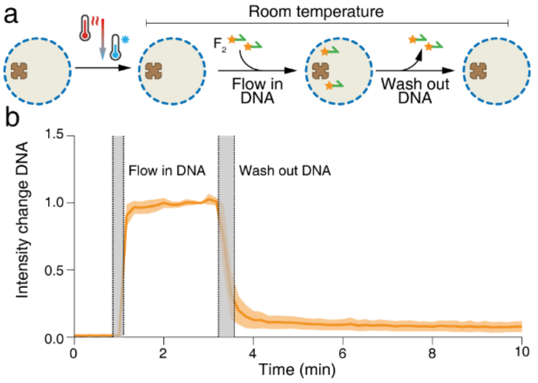

**Supplementary figure 6 Reversibility of membrane collapse. a.** Cartoon representation of experiment to determine membrane permeability after heating. Proteinosomes are heated to 95˚C for 10 minutes and subsequently cooled to room temperature before measuring membrane permeability. **b.** Normalized fluorescence intensity plot of DNA inside heated and cooled proteinosomes at room temperature over time (*n=5*). Proteinosomes were heated to 95°C for 10 minutes before cooling to room temperature and conducting the experiment. Proteinosomes were trapped in a microfluidic trapping array[1] and 50 nucleotide long Alexa546-labelled DNA $F_2$ was added. After the fluorescent signal had stabilised, buffer was flowed in to remove excess DNA. Sequence for $F_2$ is listed in Supplementary Table 1.

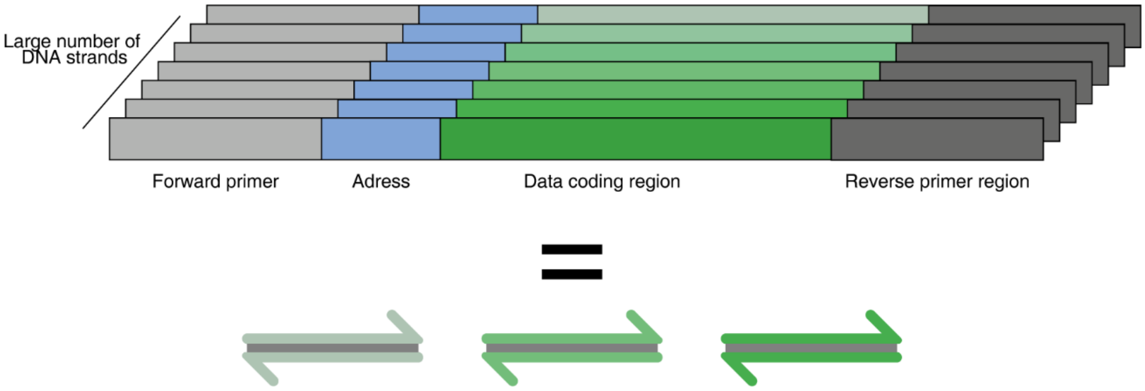

**Supplementary figure 7 Graphical representation of a general DNA file.** The procedure for the encoding a digital file starts by splitting the original data stream into appropriately sized segments. These segments are then appended with their position within the file such that the segments can be placed in the correct order after sequencing and decoding. To correct the errors inherent to DNA synthesis and sequencing error correction codes are applied to the segments to form the final segments. The final coding segments are then translated into bases in such a way that prevents repeats of identical bases. To the final DNA sequences forward and reverse primer regions are appended to allow for PCR-based amplification and retrieval. The full collection of these strands containing primer regions, address, and codign region are considered a DNA file in this work. For a detailed description of individual encoding solutions we refer the reader to references 3–5.

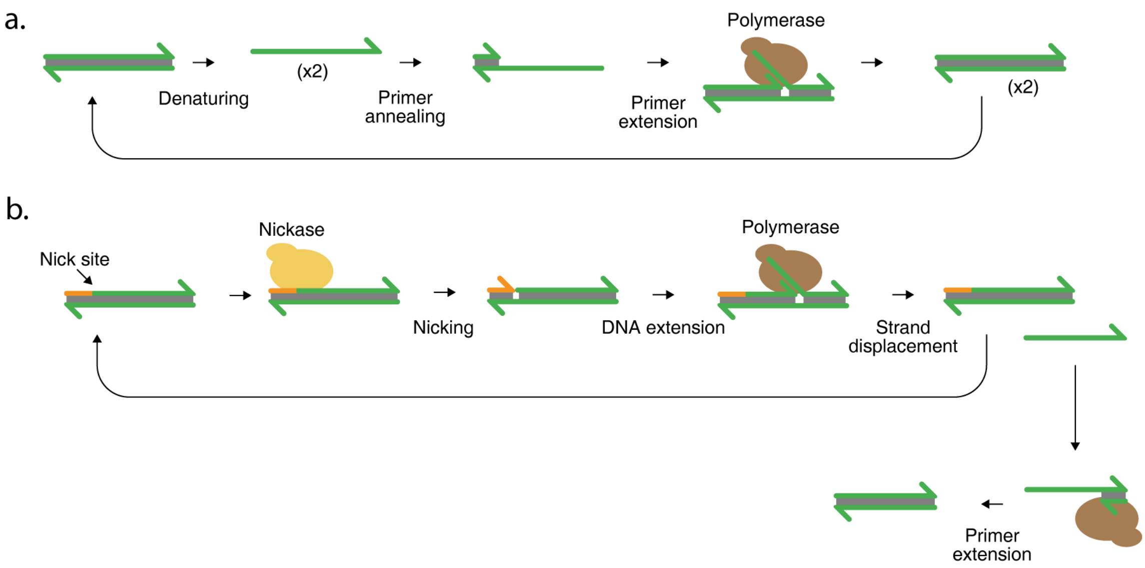

**Supplementary figure 8 Cartoons of DNA amplification methods used. a.** Schematic describing polymerase chain reaction (PCR). A piece of dsDNA is first denatured to release two single stranded DNA molecules (we draw only one of these for clarity). A short DNA primer anneals to the longer template strand and is extended by the DNA polymerase to create a dsDNA copy of the template strand. This process leads to two copies per original dsDNA template and is repeated for a fixed number of cycles. **b.** Schematic describing strand displacement amplification (SDA). A dsDNA template containing a recognition site for a nicking endonuclease (nickase) is cleaved by the nickase to generate a short DNA basepaired to the template strand and a longer secondary template. The primer is extended by a DNA polymerase with high strand displacement activity, thus also displacing the secondary template. The reformed template molecule can then again be nicked and extended to create more secondary template strands. The single stranded secondary template is converted to dsDNA using an additional primer extension reaction.

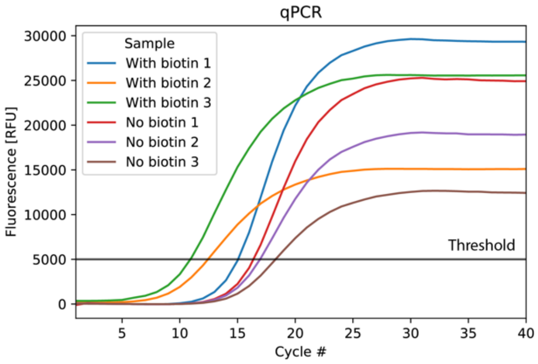

**Supplementary figure 9 Individual traces qPCR experiment summarized in Fig. 2b.** Individual traces for all triplicate qPCR experiments of proteinosomes incubated with complexes $\boldsymbol{A_1T_1}$ (with biotin) or $\boldsymbol{U_1T_{1S}}$ (without biotin). Traces were baseline adjusted using the data of the first three cycles. The threshold value was set to 5000 RFU and is plotted as a line to help the reader. Sequences for $\boldsymbol{A_1}$, $\boldsymbol{T_1}$, $\boldsymbol{U_1}$, and $\boldsymbol{T_{1S}}$ are listed in supplementary Table 2.

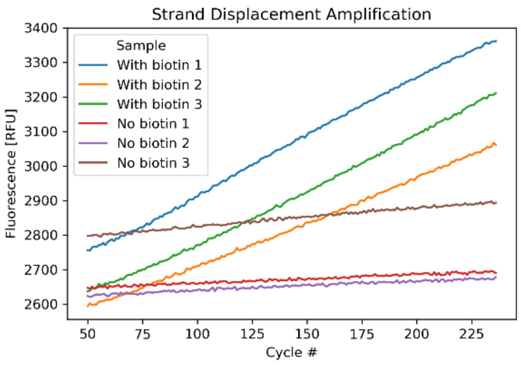

**Supplementary figure 10 Individual traces of strand displacement amplification experiment summarized in Fig. 2c.** Fluorescence versus cycle number data for individual SDA reactions of proteinosomes incubated with complexes $\boldsymbol{A_1T_1}$ (with biotin) or $\boldsymbol{U_1T_{1S}}$ (without biotin). Each cycle is approximately 30 seconds since the qPCR machine does not support exporting time-course data, we show cycle numbers. Sequences $\boldsymbol{A_1}$, $\boldsymbol{T_1}$, $\boldsymbol{U_1}$, and $\boldsymbol{T_{1S}}$ are listed in supplementary Table 2

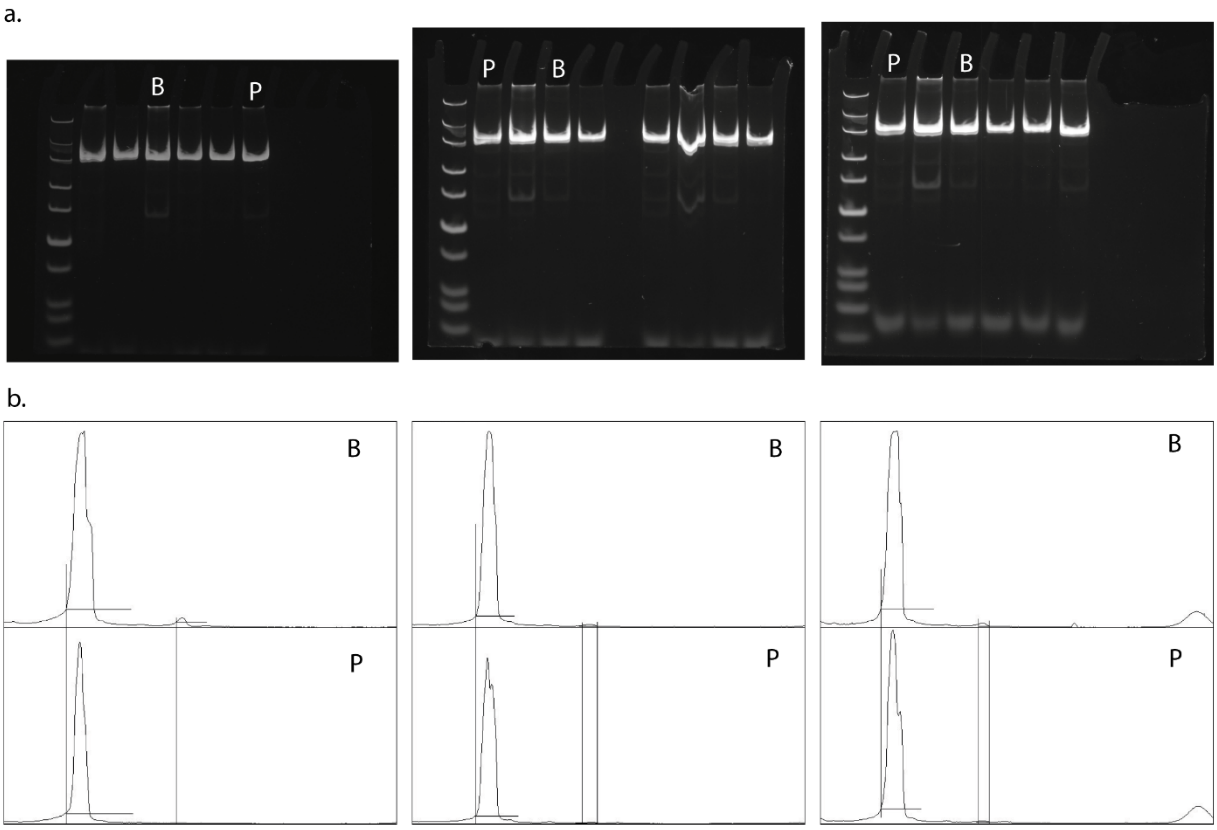

**Supplementary figure 11 Data analysis individual gels Fig. 3d. a.** Raw images of independent PAGE gels used to determine ratios of chimaera to target shown in figure 3d. B indicates bulk multiplex PCR, P indicates proteinosome localized amplified DNA. The first gel is the unprocessed image of Fig. 3c, the remaining gels are additional gels run at a later stage. Varying amounts of starting bulk DNA were used such that final concentrations match that of DNA obtained from amplification inside proteinosomes to ensure chimaera formation is not driven solely by differences in concentrations. **b.** FIJI/ImageJ gel analysis plots of lanes indicated in **a**. Lines were drawn manually to extract the area under the curves for the different products. Ratios of peaks are shown in Fig. 3d.

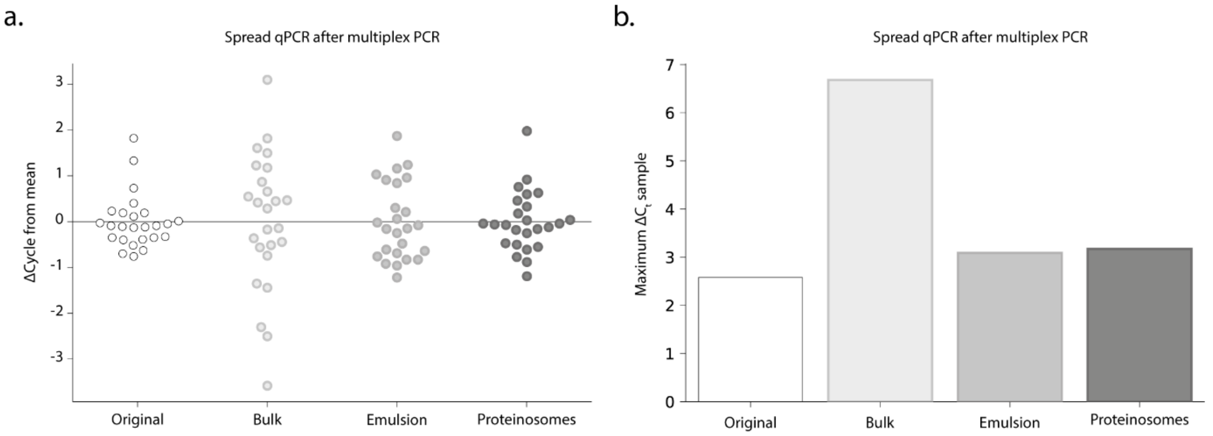

**Supplementary Figure 12 PCR inside physically separated compartments maintain the original spread of concentrations in DNA library. a.** A library consisting of 25 files encoded in DNA was amplified using either bulk PCR, thermoconfined, or emulsion PCR. These samples were compared against the distribution in the original pool and Ct-values obtained with qPCR were used as an approximation of relative concentrations. **b.** Maximum spread in concentration per condition as determined by qPCR. The lowest Ct-value was subtracted from the highest to represent the maximum fold difference in ΔCt.

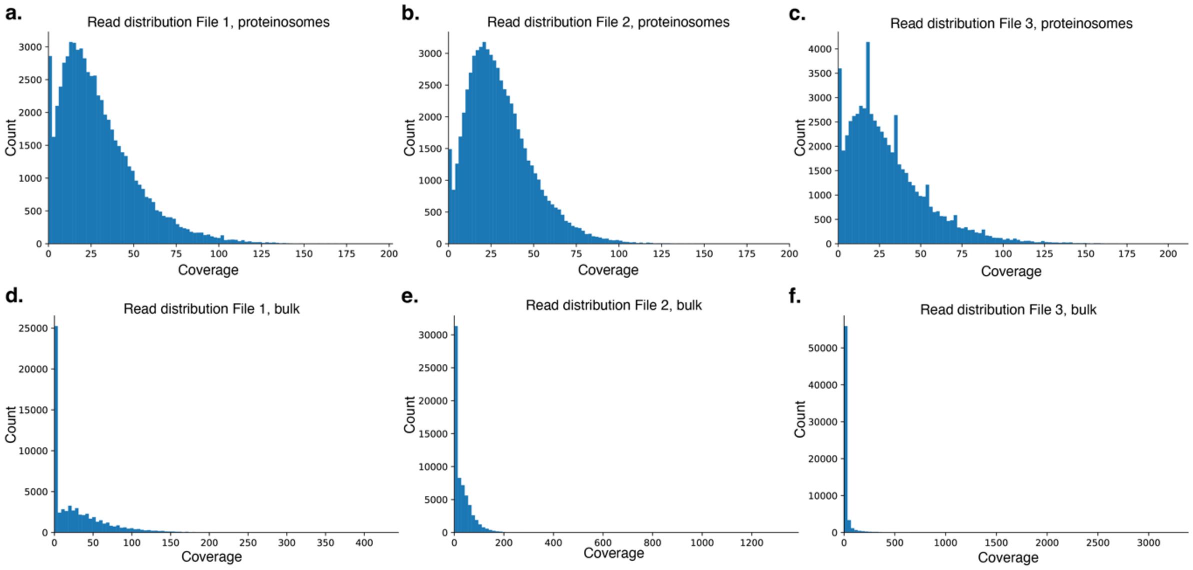

**Supplementary Figure 13 Coverage distributions after repeated PCR.** Coverages for each unique strand that mapped to a reference sequence used for encoding files are shown as determined by Illumina sequencing, randomly sampled to coverage of 30× for direct comparison. Figures **a.**, **b.** and **c.** show the sequencing distributions for proteinosome-based PCR. Figures **d.**, **e.** and **f.** show the sequencing distribution for bulk amplified DNA.

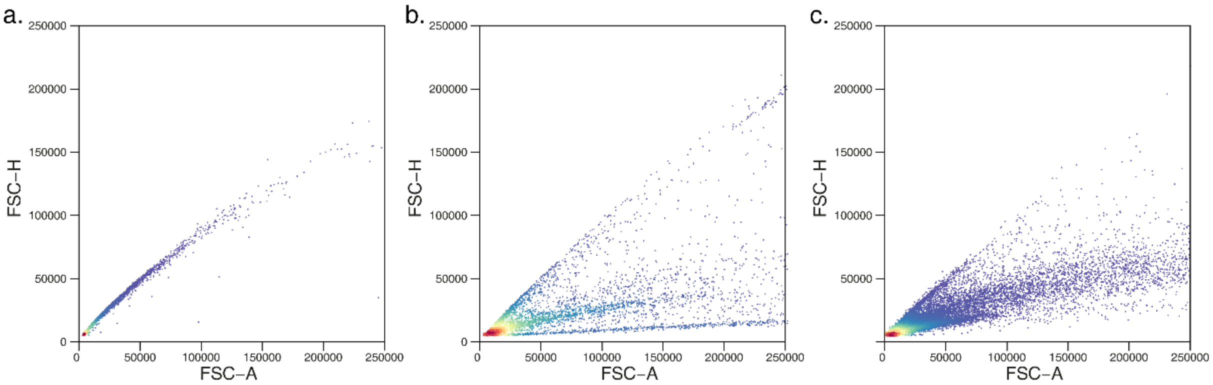

**Supplementary Figure 14 Flow cytometry analysis comparing beads and proteinosomes. a.** FSC-A versus FSC-H dotplot of magnetic particles used for preparation of proteinosomes (*n=10000*). **b.** FSC-A versus FSC-H dotplot of proteinosomes without magnetic particles (*n=5244*). **c.** FSC-A versus FSC-H dotplot of proteinosomes with magnetic particles (*n=27199*).

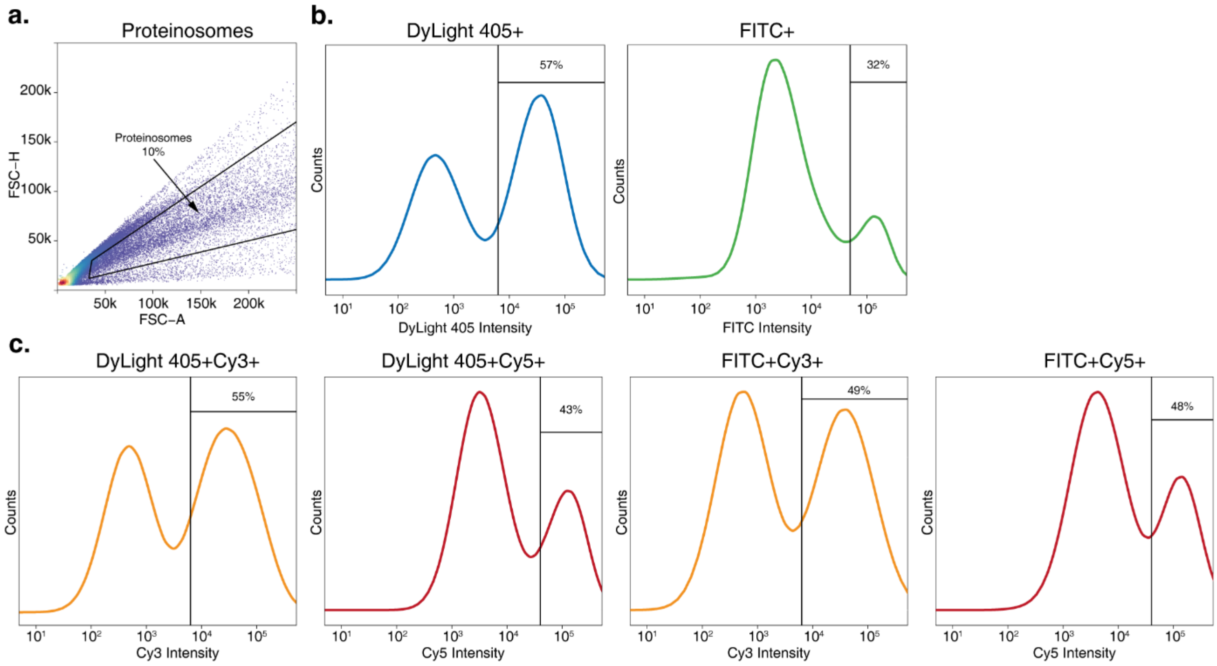

**Supplementary Figure 15 Gating strategy used to sort four proteinosome populations. a.** FSC-H vs FSC-A dotplot of initially detected events. Based on the experiments shown in Supplementary Fig. 16 a gate was drawn to select proteinosomes against background signal and unincorporated magnetic particles. Proteinosomes accounted for approximately 10% of the total number of detected events. **b.** Histograms of membrane colour intensities detected in proteinosomes gate. Gates were drawn to select high intensity populations in both DyLigh405 and FITC channels. 57% of the total proteinosome population fell within the high DyLigh405 gate, 32% of the total fell within the high FITC gate. **c.** Histograms of DNA barcode fluorescence intensities in high DyLight405 and FITC populations. Proteinosomes gated by membrane colour were subsequently gated in the Cy3 and Cy5 channels to gate based on localized barcodes. In the high DyLight405 populations 55% and 43% of the population was determined to be high Cy3 and high Cy5 respectively. In the high FITC population the high Cy3 population accounted for 49% of the total and high Cy5 for 48%.

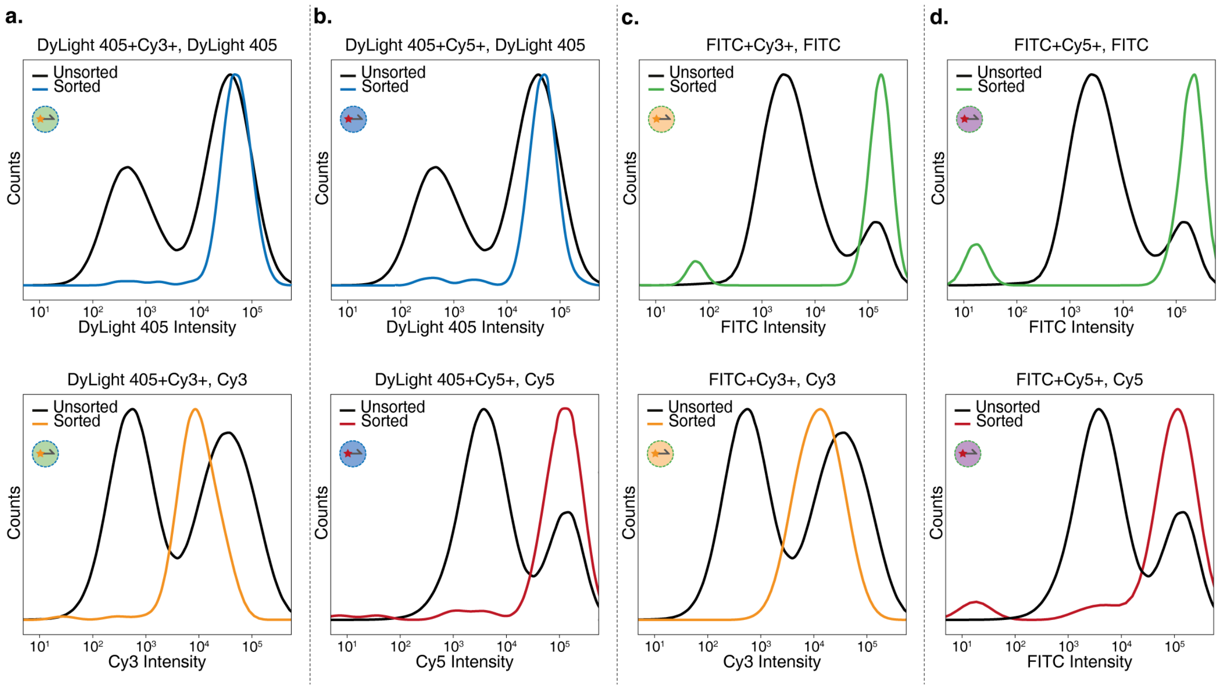

**Supplementary Figure 16 Results post-sort flow cytometry. a.** DyLight405 (blue) and Cy3 (orange) histograms fluorescence intensities of proteinosomes sorted into high DyLight405 and high Cy3 population (*n=144*). **b.** DyLight405 and Cy5 (red) histograms fluorescence intensities of proteinosomes sorted into high DyLight405 and high Cy5 population (*n=125*). **c.** FITC and Cy3 histograms fluorescence intensities of proteinosomes sorted into high FITC and high Cy3 population (*n=70*). **d.** FITC (green) and Cy5 histograms fluorescence intensities of proteinosomes sorted into high FITC and high Cy5 population (*n=81*). Single peaks are observed after sorting as opposed to the initial bimodal distributions indicating successful sorting for all four populations.

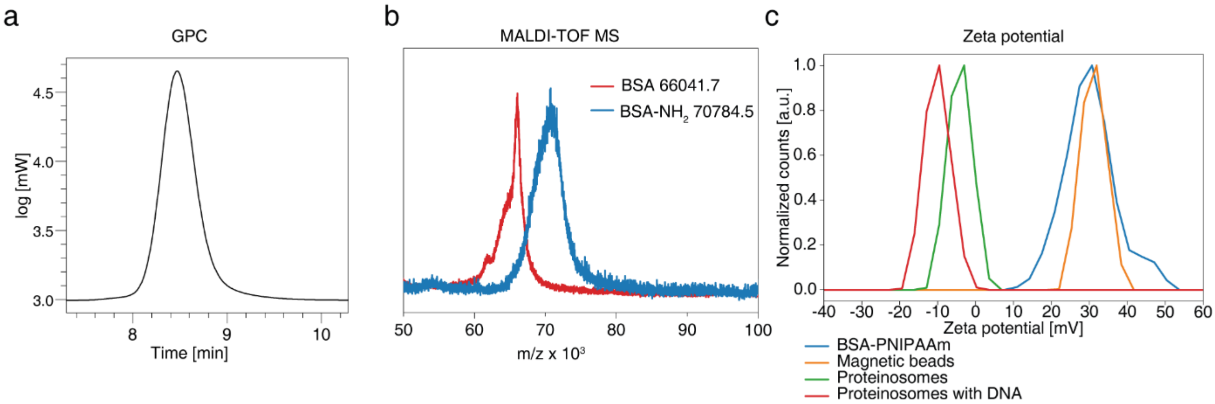

**Supplementary Figure 17 Characterization proteinosome components. a.** Gel permeation chromatograph of PNIPAAm used to prepare BSA-PNIPAAm conjugates. The molecular weight ($M_w$) was measured to be 13,284 Da, dispersity was determined to be 1.08. **b.** Matrix-assisted Laser Desorption/Ionization Time of Flight Mass Spectrometry (MALDI-TOF MS) characterization of BSA and cationized BSA. Native BSA had a molecular weight of 66,041.7 Da, after cationization the weight increased to 70,784.5 Da. **c.** Zeta potential measurements of BSA-PNIPAAm, magnetic beads, proteinosomes and DNA-loaded proteinosomes. The BSA-$NH_2$/PNIPAAm nanoconjugates have a zeta potential of 24.5 mV (Blue). Amine-modified magnetic beads have a zeta potential of 32.1 mV (orange). Empty proteinosomes have a negative zeta potential (-4.45 mV, green) because the BSA-$NH_2$ molecules in the proteinosome membrane were cross-linked resulting in a reduction in the number of surface amine groups. The charge implies that DNA does not electrostatically bind to the membrane and diffuse passively into the lumen. The zeta potential of proteinosome further decreased after localization of negatively charged DNA (-10.5 mV, red).

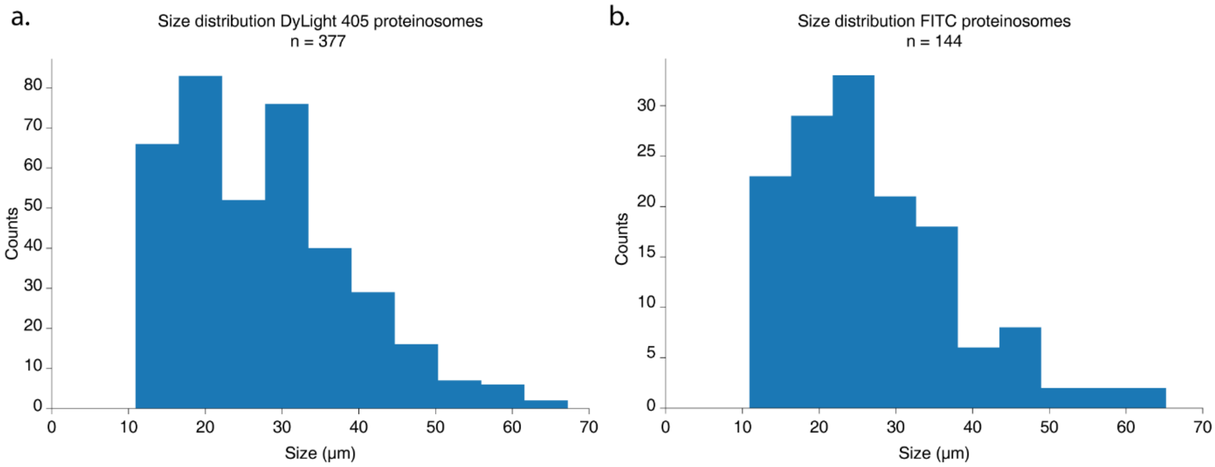

**Supplementary Figure 18 Size distribution proteinosomes used for FACS after filtering. a.** Size distribution of DyLight 405-labelled proteinosomes. The median diameter was determined to be 25.9 µm. **b.** Size distribution of FITC-labelled proteinosomes. The median diameter was determined to be 25.0 µm.

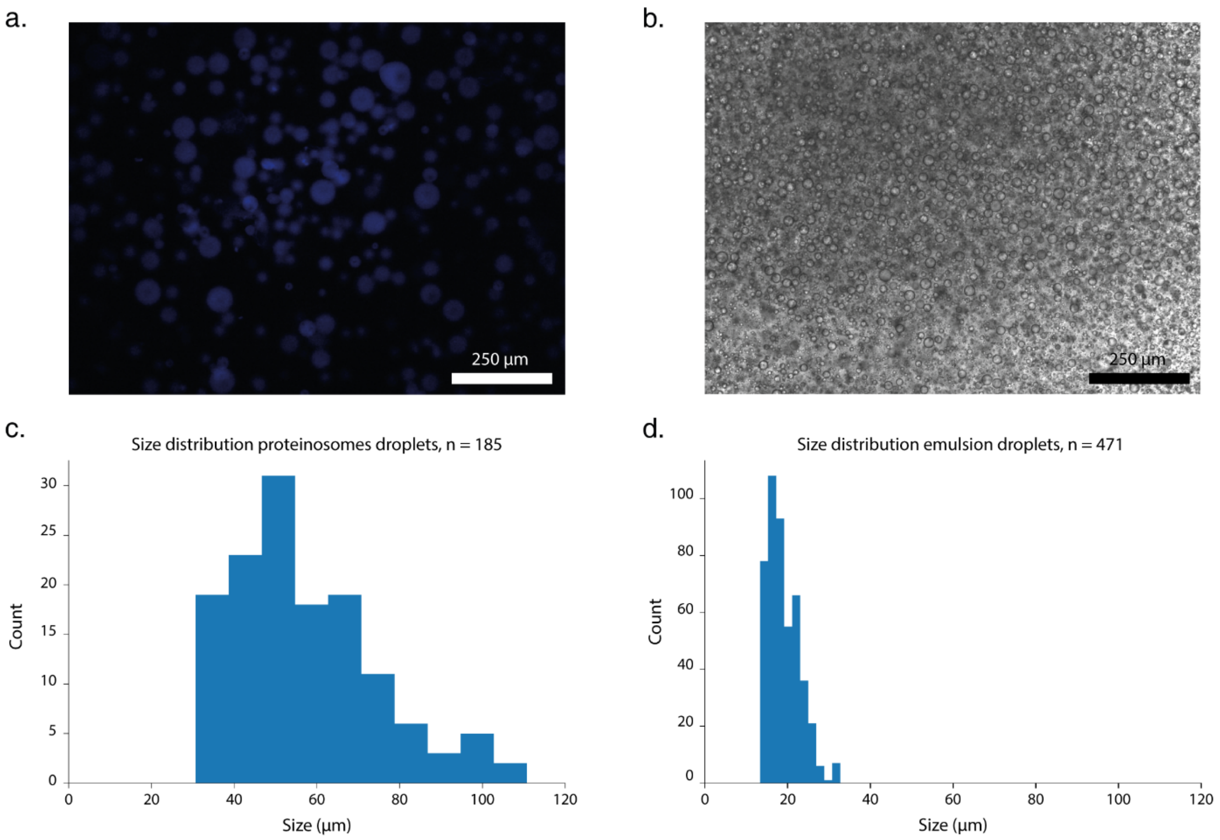

**Supplementary Figure 19 Sizes of compartments used for localized PCR. a.** Fluorescent micrograph of proteinosomes on a microscopy slide at room temperature. **b.** Brightfield micrograph of emulsion on a microscopy slide at room temperature. Emulsion PCR mixture was prepared as recommended by manufacturer, except water was used instead of PCR reaction mixture and imaged using brightfield microscopy. **c.** Size distribution of proteinosomes. The micrograph shown in **a** was analysed using FIJI/ImageJ to determine the diameter of proteinosomes and plotted in a histogram. **d.** Size distribution of emulsion droplets. The micrograph shown in **b** was analysed using FIJI/ImageJ to determine the diameter of proteinosomes and plotted in a histogram.

### Supplementary Tables

**Supplementary Table 1 – DNA sequences used in Figure 1**

| Name | Sequence | Length (# bases) | 5' modification |
|---|---|---|---|
| A1 | TTTTTTTTTTTGTCTCACTAGGTCCTCTCAGCTCTTCC CACCACCATTGCCTTCTTCTTCCTTCTGAGTCTAACCA CCACCATTGCCTTATCTACCATTCTGAGTCTTACCAC CACCATTGCCATTTCATTATCTACGAGTCTATCCACC ACCATTGCAGGAGTTTCAGTTTCGACGCAAAAAAAA AA | 188 | Biotin |
| F1 | TTTTTTTTTTGCGTCGAAACTGAAACTCCTGCAATGG TGGTGGATAGACTCGTAGATAATGAAATGGCAATG GTGGTGGTAAGACTCAGAATGGTAGATAAGGCAAT GGTGGTGGTTAGACTCAGAAGGAAGAAGAAGGCA ATGGTGGTGGGAAGAGCTGAGAGGACCTAGTGAG ACAAAAAAAAAAA | 188 | Cy5 |
| A2 | TGAGAGGACCTAGTGAGACAA | 21 | Biotin |
| F2 | TAGGTCCTCTCAGCTCTTCCCACCACCATTGCCTTCT TCTTCCTTCTGAG | 50 | Alexa546 |
| F3 | TTTTTTTTTTTTGTCTCACTAGGTCCTCTCA | 31 | Alexa546 |

**Supplementary Table 2 – DNA sequences used in Figure 2**

| Name | Sequence | Length (# bases) | 5' modification |
|---|---|---|---|
| A1 | TTTTTTTTTTTGTCTCACTAGGTCCTCTCAGCTCTTCC CACCACCATTGCCTTCTTCTTCCTTCTGAGTCTAACCA CCACCATTGCCTTATCTACCATTCTGAGTCTTACCAC CACCATTGCCATTTCATTATCTACGAGTCTATCCACC ACCATTGCAGGAGTTTCAGTTTCGACGC | 178 | Biotin |
| T1 | GCGTCGAAACTGAAACTCCTGCAATGGTGGTGGAT AGACTCGTAGATAATGAAATGGCAATGGTGGTGGT AAGACTCAGAATGGTAGATAAGGCAATGGTGGTGG TTAGACTCAGAAGGAAGAAGAAGGCAATGGTGGTG GGAAGAGCTGAGAGGACCTAGTGAGACAAAAAAA AAAA | 178 | |
| U1 | TGTCTCACTAGGTCCTCTCAGCTCTTCCCACCACCAT TGCCTTCTTCTTCCTTCTGAGTCTAACCACCACCATT GCCTTATCTACCATTCTGAGTCTTACCACCACCATTG CCATTTCATTATCTACGAGTCTATCCACCACCATTGC AGGAGTTTCAGTTTCGACGC | 168 | |

| | | |
|---|---|---|
| T1S | GCGTCGAAACTGAAACTCCTGCAATGGTGGTGGAT AGACTCGTAGATAATGAAATGGCAATGGTGGTGGT AAGACTCAGAATGGTAGATAAGGCAATGGTGGTGG TTAGACTCAGAAGGAAGAAGAAGGCAATGGTGGTG GGAAGAGCTGAGAGGACCTAGTGAGACA | 168 |
| Forward primer | TGTCTCACTAGGTCCTCTCA | 20 |
| Reverse primer | GCGTCGAAACTGAAACTCCT | 20 |

**Supplementary Table 3 – DNA sequences used in Figure 3**

| **Name** | **Sequence** | **Length (# bases)** | **5' modification** |
|---|---|---|---|
| A1 | TTTTTTTTTTGCGTCGAAACTGAAACTCCTGCAATGG TGGTGGATAGACTCGTAGATAATGAAATGGCAATG GTGGTGGTAAGACTCAGAATGGTAGATAAGGCAAT GGTGGTGGTTAGACTCAGAAGGAAGAAGAAGGCA ATGGTGGTGGGAAGAGCTGAGAGGACCTAGTGAG ACA | 178 | Biotin |
| T1 | TGTCTCACTAGGTCCTCTCAGCTCTTCCCACCACCAT TGCCTTCTTCTTCCTTCTGAGTCTAACCACCACCATT GCCTTATCTACCATTCTGAGTCTTACCACCACCATTG CCATTTCATTATCTACGAGTCTATCCACCACCATTGC AGGAGTTTCAGTTTCGACGCAAAAAAAAAA | 178 | |
| A2 | TTTTTTTTTTGCGTCGAAACTGAAACTCCTACCAATG GGGAGCCGAGGTCATCGAGTGATTACCAGCGGCTA GACTATCTAACGAGCCATTAGAGGCTGACCAGAGC AGACGTACCGCCTGAAATATGCAATGGTGGTGGAT AGACTCGTAGATAATGTGAGAGGACCTAGTGAGAC A | 178 | Biotin |
| T2 | TGTCTCACTAGGTCCTCTCACATTATCTACGAGTCTA TCCACCACCATTGCATATTTCAGGCGGTACGTCTGCT CTGGTCAGCCTCTAATGGCTCGTTAGATAGTCTAGC CGCTGGTAATCACTCGATGACCTCGGCTCCCCATTG GTAGGAGTTTCAGTTTCGACGCAAAAAAAAAA | 178 | |
| C1 | TGTCTCACTAGGTCCTCTCACATTATCTACGAGT CTATCCACCACCATTGCAGGAGTTTCAGTTTCG ACGC | 71 | |
| C2 | GCGTCGAAACTGAAACTCCTGCAATGGTGGTG GATAGACTCGTAGATAATGTGAGAGGACCTAG TGAGACA | 71 | |
| Forward primer | TGTCTCACTAGGTCCTCTCA | 20 | |
| Reverse primer | GCGTCGAAACTGAAACTCCT | 20 | |

**Supplementary Table 4 – DNA sequences used in Figure 4**

| Name | Sequence | Length (# bases) | 5' modification |
|---|---|---|---|
| File 1 Forward | AAAGCGTCGGAAGTTTGTGA | 20 | |
| File 1 Reverse | TTTCGCGGTGGTTGTTATGT | 20 | |
| File 2 Forward | AACTGCTTAATCGTGCCACA | 20 | |
| File 2 Reverse | ACAAACTTCGAGAACTGCGT | 20 | |
| File 3 Forward | ATGAGTGCGTCTTGTGTGTT | 20 | |
| File 3 Reverse | ACCGCTTTGACGCAAACTAA | 20 | |
| File 4 Forward | TTAATCGGTAACACCTGCGG | 20 | |
| File 4 Reverse | CACTTTAGCCTAACGGTGGT | 20 | |
| File 5 Forward | ACTGCCAACGTATTGCACAT | 20 | |
| File 5 Reverse | AAACCAGACCGTTGTCGAAA | 20 | |
| File 6 Forward | AATTTGGCATTCACCGTGGA | 20 | |
| File 6 Reverse | TTTCATTGGCTTGCACCAGA | 20 | |
| File 7 Forward | TCACGACTTTGTTTGCGAGT | 20 | |
| File 7 Reverse | AAGTTAGTTCAACGTCGCGT | 20 | |
| File 8 Forward | TTGGTTGACCGTGTGTGAAT | 20 | |
| File 8 Reverse | ATAACGCGCAAGAAGCATCT | 20 | |
| File 9 Forward | AATGAAGATGCACGCCTTGA | 20 | |
| File 9 Reverse | ACAAGCATTCCGTCACAGTT | 20 | |
| File 10 Forward | AATGGACGTTCCGCAATCAT | 20 | |
| File 10 Reverse | ATTAGCCAAACCATAGCGCA | 20 | |
| File 11 Forward | TTACCGTGCAGTTGACGAAA | 20 | |
| File 11 Reverse | ATTGTTACGAATCGGTGCCA | 20 | |
| File 12 Forward | ATCTGCGCTTAACAAGGCTT | 20 | |
| File 12 Reverse | AGTCGCCAAATAAGTGCCAT | 20 | |
| File 13 Forward | TGGTTTCGTGTTCAAGCGTA | 20 | |
| File 13 Reverse | CGCCATTGCAGAAAGAGAGA | 20 | |
| File 14 Forward | AAACAAAGTTAGCGGCTCGT | 20 | |
| File 14 Reverse | AATAACCGCGAATTGGCCTT | 20 | |
| File 15 Forward | TGGTTTGCAGGAAATGTCGT | 20 | |
| File 15 Reverse | ACCGCTTTGCACTCAGTTAA | 20 | |
| File 16 Forward | ACCGCGCTCGAAGAATTTAA | 20 | |
| File 16 Reverse | TACAGGCAACAAACGCATCT | 20 | |
| File 17 Forward | TCTCCGGTGCTACGTTAAAG | 20 | |
| File 17 Reverse | CGGCCTTCTCCAAAGAATCA | 20 | |
| File 18 Forward | AACGAATCGAACTGCCTTGT | 20 | |
| File 18 Reverse | TGAAACATATTCGCACGCCT | 20 | |
| File 19 Forward | TCCTGCTTGCGTTAAATGGA | 20 | |
| File 19 Reverse | AAACCGACGGAAACATTGCA | 20 | |
| File 20 Forward | AACCGCTTTGGCTCAGTATT | 20 | |
| File 20 Reverse | TAAAGGCAGGCAAACGGATT | 20 | |
| File 21 Forward | AGTGCATTCTCCAAGCAACT | 20 | |

| | | | |
|---|---|---|---|
| File 21 Reverse | TACACACGGTTTGGCTTGAA | 20 | |
| File 22 Forward | TTCGCAGGTGACTTTGTGTT | 20 | |
| File 22 Reverse | ATGAGGCAAATCGTCGCAAA | 20 | |
| File 23 Forward | AACCGATGCGCTTATACGTT | 20 | |
| File 23 Reverse | AGGAAGCGCCAACTAATTGT | 20 | |
| File 24 Forward | AGCCTTGTGTCCATCAATCC | 20 | |
| File 24 Reverse | TTGTTCAGTGGAAGACGACG | 20 | |
| File 25 Forward | AAGGCTCGCGTGCATATTAA | 20 | |
| File 25 Reverse | ATTATGAAAGCGCAGGCAGA | 20 | |
| File 1 Forward bio | TTTTTTTTTTTAAAGCGTCGGAAGTTTGTGA | 31 | Biotin |
| File 2 Forward bio | TTTTTTTTTTTAACTGCTTAATCGTGCCACA | 31 | Biotin |
| File 3 Forward bio | TTTTTTTTTTTATGAGTGCGTCTTGTGTGTT | 31 | Biotin |
| File 4 Forward bio | TTTTTTTTTTTTTAATCGGTAACACCTGCGG | 31 | Biotin |
| File 5 Forward bio | TTTTTTTTTTTACTGCCAACGTATTGCACAT | 31 | Biotin |
| File 6 Forward bio | TTTTTTTTTTTAATTTGGCATTCACCGTGGA | 31 | Biotin |
| File 7 Forward bio | TTTTTTTTTTTTCACGACTTTGTTTGCGAGT | 31 | Biotin |
| File 8 Forward bio | TTTTTTTTTTTTTGGTTGACCGTGTGTGAAT | 31 | Biotin |
| File 9 Forward bio | TTTTTTTTTTTAATGAAGATGCACGCCTTGA | 31 | Biotin |
| File 10 Forward bio | TTTTTTTTTTTAATGGACGTTCCGCAATCAT | 31 | Biotin |
| File 11 Forward bio | TTTTTTTTTTTTTACCGTGCAGTTGACGAAA | 31 | Biotin |
| File 12 Forward bio | TTTTTTTTTTTATCTGCGCTTAACAAGGCTT | 31 | Biotin |
| File 13 Forward bio | TTTTTTTTTTTTGGTTTCGTGTTCAAGCGTA | 31 | Biotin |
| File 14 Forward bio | TTTTTTTTTTTAAACAAAGTTAGCGGCTCGT | 31 | Biotin |
| File 15 Forward bio | TTTTTTTTTTTTGGTTTGCAGGAAATGTCGT | 31 | Biotin |
| File 16 Forward bio | TTTTTTTTTTTACCGCGCTCGAAGAATTTAA | 31 | Biotin |
| File 17 Forward bio | TTTTTTTTTTTTCTCCGGTGCTACGTTAAAG | 31 | Biotin |
| File 18 Forward bio | TTTTTTTTTTTAACGAATCGAACTGCCTTGT | 31 | Biotin |
| File 19 Forward bio | TTTTTTTTTTTTCCTGCTTGCGTTAAATGGA | 31 | Biotin |
| File 20 Forward bio | TTTTTTTTTTTAACCGCTTTGGCTCAGTATT | 31 | Biotin |
| File 21 Forward bio | TTTTTTTTTTTAGTGCATTCTCCAAGCAACT | 31 | Biotin |
| File 22 Forward bio | TTTTTTTTTTTTTCGCAGGTGACTTTGTGTT | 31 | Biotin |
| File 23 Forward bio | TTTTTTTTTTTAACCGATGCGCTTATACGTT | 31 | Biotin |
| File 24 Forward bio | TTTTTTTTTTTAGCCTTGTGTCCATCAATCC | 31 | Biotin |
| File 25 Forward bio | TTTTTTTTTTTAAGGCTCGCGTGCATATTAA | 31 | Biotin |

**Supplementary Table 5 – Coverage per file for each condition**

| File | Original | Bulk | Emulsion | Proteinosomes |
|---|---|---|---|---|
| 1 | 16.23 | 5.50 | 27.76 | 8.77 |
| 2 | 20.63 | 14.94 | 32.17 | 13.82 |
| 3 | 20.85 | 12.19 | 32.49 | 12.08 |
| 4 | 23.70 | 15.04 | 29.00 | 31.80 |
| 5 | 26.88 | 21.53 | 36.11 | 18.56 |
| 6 | 17.61 | 199.91 | 73.12 | 13.30 |
| 7 | 16.54 | 84.48 | 44.78 | 12.89 |
| 8 | 9.82 | 33.29 | 15.29 | 5.84 |
| 9 | 30.69 | 46.80 | 64.38 | 38.47 |
| 10 | 21.47 | 11.21 | 31.92 | 21.15 |
| 11 | 25.61 | 78.26 | 57.12 | 26.19 |
| 12 | 24.35 | 13.04 | 37.15 | 17.57 |
| 13 | 15.83 | 25.65 | 28.33 | 12.84 |
| 14 | 18.09 | 11.20 | 30.17 | 9.28 |
| 15 | 24.81 | 5.22 | 22.66 | 26.77 |
| 16 | 18.83 | 16.58 | 37.76 | 11.97 |
| 17 | 15.89 | 16.81 | 34.17 | 5.42 |
| 18 | 26.40 | 14.19 | 45.95 | 20.08 |
| 19 | 15.28 | 19.03 | 36.72 | 8.10 |
| 20 | 19.21 | 3.31 | 21.36 | 18.41 |
| 21 | 17.78 | 19.98 | 37.87 | 9.39 |
| 22 | 22.16 | 7.15 | 33.21 | 6.06 |
| 23 | 23.44 | 31.91 | 44.18 | 25.13 |
| 24 | 16.42 | 17.45 | 32.13 | 15.49 |
| 25 | 27.79 | 9.58 | 34.36 | 17.57 |

**Supplementary Table 6 – Sequences used in figure 5**

| Name | Sequence | Length (# bases) | 5' modification | 3' modification |
|---|---|---|---|---|
| F4 | GACAGGAAAGGACTCGGTGGGT TAGGCAATGGATCAG | 37 | Biotin | Cy3 |
| F5 | GTTAGACTCTTAGGGTGGTGC AATGGGGTAT | 31 | Biotin | Cy5 |
| File 1 Forward | TGGTTTCGTGTTCAAGCGTA | 20 | | |
| File 1 Reverse | CGCCATTGCAGAAAGAGAGA | 20 | | |
| File 2 Forward | AAACAAAGTTAGCGGCTCGT | 20 | | |
| File 2 Reverse | AATAACCGCGAATTGGCCTT | 20 | | |
| File 3 Forward | TGGTTTGCAGGAAATGTCGT | 20 | | |
| File 3 Reverse | ACCGCTTTGCACTCAGTTAA | 20 | | |

| | | |
|---|---|---|
| File 4 Forward | ACCGCGCTCGAAGAATTTAA | 20 |
| File 4 Reverse | TACAGGCAACAAACGCATCT | 20 |
